# Noise Distorts the Epigenetic Landscape and Shapes Cell Fate Decisions

**DOI:** 10.1101/2020.12.21.423724

**Authors:** Megan A. Coomer, Lucy Ham, Michael P. H. Stumpf

## Abstract

The Waddington epigenetic landscape has become an iconic representation of the cellular differentiation process. Recent single-cell transcriptomic data provide new opportunities for quantifying this originally conceptual tool, offering insight into the gene regulatory networks underlying cellular development. While many methods for constructing the landscape have been proposed, by far the most commonly employed approach is based on computing the landscape as the negative logarithm of the steady-state probability distribution. Here, we use simple models to highlight the complexities and limitations that arise when reconstructing the potential landscape in the presence of stochastic fluctuations. We consider how the landscape changes in accordance with different stochastic systems, and show that it is the subtle interplay between the deterministic and stochastic components of the system that ultimately shapes the landscape. We further discuss how the presence of noise has important implications for the identifiability of the regulatory dynamics from experimental data.

## Introduction

Waddington’s epigenetic landscape has become more than just a metaphor for cellular differentiation and developmental biology. In recent years it has garnered renewed interest due to (i) technological advances in e.g. single-cell biology; and (ii) because of its close, intimate link to mathematical models that describe developmental processes in biology. Due to the excellent expository work of [1] and [2], but also through Waddington’s own writing and influence [3], the Waddington (also known as *epigenetic*) landscape, has made mathematical concepts palatable to traditionally non-mathematical audiences. The intuitive representation of a dynamical system provided by the landscape also appeals to mathematical scientists, and has done so for some time. René Thom was very interested in developmental biology [4] and communicated extensively with Waddington.

One of the appeals of the landscape is that it offers a useful way of making mathematical concepts accessible. Traditionally, however, the analysis (and interpretation) of Waddington’s epigenetic landscape has revolved around considering deterministic dynamics. At the cellular level, however, noise – caused by the stochasticity underlying the molecular processes involved in gene regulation, protein turnover, etc. – is ubiquitous. Noise interacts with dynamics in often non-intuitive ways [5], and it is important to set out the conditions under which noisy and deterministic dynamics agree with respect to important qualitative (and quantitative) aspects of the dynamics.

Linking the landscape to gradient systems, deterministic dynamical systems of the type

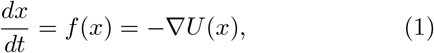

where *x* denotes the state of the system, and where *U*(*x*) is a potential function, have proven to be particularly fruitful. For such systems we can access powerful mathematical frameworks, such as Morse theory (every gradient system is a Morse function) [6] (Section 9.9), and [7] (Section 8.2 & 8.3); and a slew of further mathematical developments following Morse’s original work (for a general overview see [8, 9]). However, for systems found in biology, which are typically high-dimensional and non-equilibrium systems, the deterministic force *f*(*x*) cannot always be quantified as a pure gradient system. On the face of it, restricting our attention to gradient systems may therefore appear limiting. But there are two important results, that mean that this is in fact not the case: (i) gradient systems (or more generally, Morse-Smale systems) are very common, and any function chosen at random is a Morse function with high probability [6] (Section 9.9); (ii) systems that are not gradient systems can be transformed into gradient systems in well understood ways, see [10] for an overview. Thus gradient systems are (qualitatively and certain aspects at least semi-quantitatively) representative of more general dynamical systems [8] (Page 755).

Noisy biological systems are frequently modelled in terms of stochastic differential equations (SDEs) [11]. Here the state of the system *x*, usually defined in terms of molecular concentrations, is described by

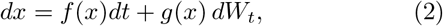

where *f*(*x*) represents the deterministic dynamics of the system and *g*(*x*)*dW*_*t*_ the stochastic dynamics, with *dW*_*t*_ an increment in the Wiener process. The components *f*(*x*) and *g*(*x*) are commonly referred to as the *drift* and *diffusion* components of the system, respectively. Often the noise component *g*(*x*) is taken to be independent of the state of the system (i.e. *g*(*x*) is constant), and has become known as *additive noise*. When the noise component *g*(*x*) depends on the state of the system, the noise is said to be *multiplicative* [12].

The recent availability of single-cell transcriptomic data has motivated a number of different approximations to potential functions that can serve as mathematical descriptions of the epigenetic landscape [13, 14, 15]. Such approximations are referred to as *quasi-potential landscapes* and are realised using different, yet often overlapping, methods that rely on computing a potential or quasi-potential based on the SDE given in Eq. (2) above. In general, the drift component might not arise from the gradient of a potential function, and therefore cannot be described by Eq. (1) alone. Rather, the deterministic component *f*(*x*) may be decomposed into the sum of the gradient of a potential, *U*(*x*), and a remaining non-gradient component referred to as the *curl flux* [16].

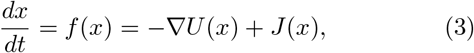

The gradient portion of the dynamics determines the most probable states of the dynamical system (and thus the basins of attraction in the potential landscape), while the curl flux determines the paths traversed between them; it provides an additional rotational contribution to the motion, over and above the gradient, and can result in divergence from the paths of steepest descent (or gradient paths) in the system [17, 18]. The potential therefore provides an intuitive description of the developmental landscape: the system will tend to stay near to regions of low potential (the minima), and will move away from regions with high potential (the maxima). Numerous approaches for determining the decomposition of the SDE in which the potential is defined have been proposed. Examples include the Helmholtz decomposition [19], the normal or orthogonal decomposition [13], the symmetric-anti-symmetric decomposition [20], the decomposition based on the probability flux in steady-state conditions [16], and the so-called Freidlin-Wentzell quasi-potential formulated on large deviation theory for stochastic processes [21]. Each decomposition method realises a different quasi-potential function with its own unique advantages and disadvantages. These decomposition methods, and importantly how they formally relate to one another, is well-reviewed in [13, 14]. The choice of decomposition largely depends on the biological question at hand, but by far the most popular technique for reconstructing the potential landscape is based upon the steady-state probability distribution, *P*_*s*_(*x*), of the stochastic system given in Eq (2) [16, 22, 23, 24, 25, 26, 27]. Here the elevation of the landscape relates to the inverse of the probability density function of the gene expression space. States with the highest probability density, indicated by peaks in the distribution, will have the lowest potential, and thus will correspond to valleys on the landscape. More precisely, the approximated potential landscape is evaluated as the Boltzmann-like distribution from equilibrium statistical mechanics [16],

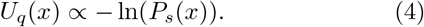

In practise, the steady-state distribution is obtained in one of two ways: (1) directly from experimental data, or (2) by solving the associated Fokker-Planck equation (FPE), either analytically [28, 29, 30] or, more commonly, by way of simulations of the corresponding SDE [16, 22, 31, 26, 32, 27]. To avoid ambiguity, throughout we refer to the estimated landscape *U*_*q*_(*x*) as the *probabilistic landscape*, and refer to *U*(*x*) as the *potential* or *deterministic landscape*.

While the steady-state distribution obtained directly from experimental data has proven successful in some applications [33], the most fruitful approach is supported by a combination of both experimental and *in silico* methods. A common approach is to first construct a core regulatory model using the support of experimental data, from which an SDE is derived. The associated FPE is then solved in steady-state conditions by way of (typically) extensive simulations of the SDE. The quasi-potential landscape is computed via the steady-state probability distribution according to Eq. (4) above. The topography of the landscape may then provide insight into the different attractor states or phenotypes and the relative stabilities between them, facilitating the mechanistic understanding of the underlying process [34]. Moreover, kinetic transition paths between stable states can give us an intuition into differentiation, de-differentiation and lineage reprogramming [35].

In this paper, we consider simple one-dimensional stochastic systems to examine the recovery of the potential landscape *U*(*x*) by way of the probabilistic landscape *U*_*q*_(*x*), using Eq. (4). We demonstrate mathematically that this recovery is strongly dependent on interplay between noise and the deterministic dynamics of the system, as well as highlight some additional complexities that arise in the presence of biological noise. Our results are demonstrated on two model systems: (1) a one-dimensional double-well potential where the drift and diffusion components are independent, and (2) a simple chemical reaction network model, where the diffusion term is coupled to the underlying deterministic dynamics. The first model is reminiscent of those often considered in the study of developmental systems. In these studies, it is assumed that the number of molecules in the system is large enough that intrinsic fluctuations may be neglected, and the effects of an independent extrinsic noise term on the underlying deterministic dynamics is considered. Our analysis shows how the inclusion of an external noise term can have a significant impact on the dynamics of the system, resulting in qualitative differences between the deterministic and probabilistic landscapes. In particular, we show how multiplicative noise is capable of shifting, creating or even destroying fixed points in the landscape. Our second model, which is based upon a set of simple chemical reactions, is then used to demonstrate how noise arising from internal fluctuations alone can also have a significant impact on the system dynamics, and subsequently the probabilistic landscape. We use our model to highlight some of the limitations in common modelling practices adopted in the study of developmental systems.

Beyond the equilibrium case we consider here, there exist a number of computational frameworks to distill cellular trajectories and landscapes from single-cell data along a time-course [36, 37, 38, 39]. Irrespective of the temporal setting however, all of these reconstruction methods face a common challenge: transcriptional profiling allows measurements at only single instances (snapshots), and cannot provide the temporal variation of any individual cell. This places fundamental limits on the identifiability of the system dynamics [26, 40]. Recent developments that leverage multimodal and high-dimensional experimental techniques [40, 41, 42] offer some progress towards the full identification of regulatory dynamics, but still more work needs to be done.

Finally, we use a simple stochastic system to illustrate some of the limitations to inferring the underlying regulatory dynamics of a noisy system from experimental data. More specifically, we show that knowledge of the typical distribution of cell states alone, as given by the steady-state probability distribution, is not sufficient to make definitive assumptions about the underlying dynamics.

The systems we consider display rich qualitative behaviour, but are simple enough to be analytically tractable. Throughout, we discuss how our results relate to real, typically higher-dimensional, biological systems.

### Noisy gene regulatory networks shape the landscape

During differentiation, we often observe increased variability in the expression of specific genes between cells of the same type [43, 44, 45, 46, 47]. These changes in transcriptional variability are intrinsically linked to the inherent stochasticity of the transcription and translation processes [48, 49, 50, 51], and appear to be intimately related to differentiation initiation and commitment [52]. As such, the epigenetic landscape, shaped by the underlying gene regulatory network (GRN), is subject to the effects of stochastic fluctuations or noise.

Noise may impact the landscape differently depending on the deterministic and stochastic dynamics of the system. For example, noise may have a purely disorganising effect on the landscape, where the deterministic dynamical structure remains largely intact (i.e. the position and number of valleys and hills on the landscape remain fixed), but cells jostle around the system and occasionally, with an increase in noise intensity, hop from one stable state to another [12]. Mathematically, this changes the steady-state distribution *P*_*s*_(*x*) from a family of delta-peaks to a broader distribution, with mass centered around the family of peaks – often referred to as a “flattening” of the distribution. By way of Eq. (4), this also corresponds to a flattening of the landscape. In addition to these effects, noise may result in qualitative changes to the landscape, by modifying both the position *and* number of valleys and hills [12]. Here the mathematical equivalent is not only a broadening of *P*_*s*_(*x*), but a shift in positions of the mode(s), as well as the appearance or disappearance of modes. We will see below how noise can drive the creation of new valleys and hills, or destroy existing ones on the deterministic landscape.

#### Box 1 Notation and Terminology

*x* — Variable denoting the state of the system

*f*(*x*) — Deterministic dynamics of a stochastic dynamical system

*g*(*x*)*dWt* — Stochastic dynamics of a stochastic dynamical system

*dWt* — Increment in the Wiener process

*P*_*s*_(*x*) — Steady-state probability distribution of a stochastic dynamical system

*U*(*x*) — Potential function or *deterministic landscape*

*U*_*q*_(*x*) — *Probabilistic landscape* computed as logarithm of the inverse of the steady-state probability distribution of a stochastic system

*α* — Stochastic integration convention

*σ* — Noise intensity

*σ*_*c,α*_ — Critical noise (resulting in a qualitative change in the steady-state probability distribution) for a given stochastic integration convention

Stochastic fluctuations inherent to gene expression arise from both internal and external sources [48, 53, 54, 55, 56] (otherwise known as *intrinsic* or *extrinsic* noise, respectively). The former originates from probabilistic chemical reactions taking place inside the cell at low concentrations, while the latter refers to stochasticity in the cells’ ever-changing physical and chemical environment. Consequently, the stochastic dynamics of any GRN reflect both the internal and external noise sources at play. Stochasticity arising from internal fluctuations is most accurately described by the Chemical Master Equation (CME). The CME is however, notoriously difficult to solve with only a handful of simple systems admitting an analytical solution. As a consequence, a number of different methods for simulating the stochastic dynamics of GRNs have been developed, and are now routinely employed [57, 58, 59]. Gillespie’s stochastic simulation algorithm (SSA) [58] is an exact method, producing sample paths of the systems trajectory whose probability distribution is the solution of the CME. The main disadvantage of this approach is that it quickly becomes infeasible for simulating realistic biological systems, which typically involve numerous reactions, and may exhibit multiple reaction timescales. Continuous approximations such as the Chemical Langevin Equation (CLE) [59], which generate approximate sample paths, can afford significant computational advantages, and have been widely adopted as alternative frameworks for simulating the stochastic dynamics of chemical reaction systems. In the CLE framework, the noise term (corresponding to *g*(*x*) in Eq. (2)) represents only intrinsic noise, and is determined solely by the deterministic dynamics [59]. In contrast, there is no commonly accepted approach for modelling extrinsic noise. While numerous SSA-like approaches for simulating extrinsic noise have emerged [60, 61, 62, 63], there is no standard approximative approach. Often an extrinsic noise term is incorporated into Langevin-type equations by way of *ad hoc*, and sometimes physically unfounded means; the choice of noise is typically taken to be additive, which has no real physical basis. As these approaches can be justified primarily on the grounds of simplifying the properties of the system, and hence the analysis, the validity of such approaches for probing the stochastic dynamics of biological systems is unclear. In future work, we follow Gillespie’s derivation of the CLE (or alternatively, the Kramers-Moyal expansion of the CME truncated to the first two terms) to obtain an analogous approximation for systems subject to extrinsic noise arising from time-dependent rate constants. The advantage of our approach is that it is computationally efficient, and does not arise from an *ad hoc* strategy: extrinsic fluctuations are introduced at the level of the CME, and the corresponding CLE approximation is derived from this. The approach may be useful in the effort towards the reconstruction of potential landscapes, which in many biological settings, require simulating the dynamics of both intrinsic and extrinsic noise.

Furthermore, noise may enter the system in an additive or multiplicative manner. The GRN may be regulated by the internal state of the cell (e.g. stochastic fluctuations in mRNA and protein levels, feedback loops [64] and higher order interactions and dependencies [65]), as well as the cell’s microenvironment [66, 67]. Based on this, it is expected that the stochastic dynamics of the GRN are multiplicative in nature [68]. Mathematically then, noise is no longer a constant term in Eq. (2), but rather a function of the system’s state. Moreover, noise may now take on a variety of forms depending on how it relates to the state of the system. When these mathematical subtleties are taken into account, there may be substantial deviations between the deterministic and probabilistic landscapes, which we demonstrate and discuss more formally below. A natural question then arises: what form of noise is relevant for cellular differentiation, and how does it affect the landscape? Here we are guided by a growing body of experimental evidence from single-cell expression analyses, that show that cells undergoing differentiation display an increase in noise intensity at cell-fate decision points. For a recent summary of this phenomenon in different cell types see [69]. Indeed, in one of the first articles to mathematically formalise the link between stochasticity and the epigenetic landscape, the authors propose that noise is highest during cell-fate decision points of development [67].

Taken together, we consider different multiplicative noise forms that are consistent with experimentally observed cell-to-cell variability. Specifically, we consider noise where the intensity is high at cell-fate decision points, and comparatively lower at differentiated cell states^2^. We also find it pertinent to consider additional noise forms that will assist in demonstrating the nuances that arise in computing the probabilistic landscape in the presence of stochasticity.

### Multiplicative noise and reconstructing the potential landscape

As discussed above, multiplicative noise is the expectation rather than the exception in biological systems. The introduction of a state-dependent noise component, *g*(*x*), introduces a number of subtleties in using the distribution-based method given in Eq. (4) for reconstructing the potential landscape. While some of these complications are widely understood in physics [12, 70], they are perhaps less so in a biological context.

The first subtlety arises in solving the SDE in Eq. (2). The integration of Eq. (2) now involves stochastic integrals, which can be interpreted according to different conventions. Such integrals are calculated in the standard way via discretisation

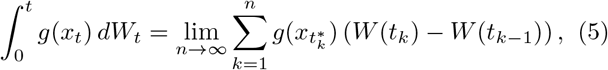

for increasingly finer partitions 0 = *t*_0_ < *t*_1_ < ... < *t*_*n*_ − 1 < *t*_*n*_ = *t*. Unlike standard integration however, the integral now depends on the point 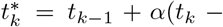 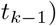 in [*t*_*k*−1_, *t*_*k*_] where the process *W*(*t*) is evaluated. Here *α* ∈ [0, 1] and is known as the *stochastic integration convention*. The two most common conventions are the Itô interpretation, where 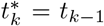 (that is, *α* = 0), and the Stratonovich interpretation, where 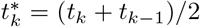 (that is, *α* = 0.5). Allowing for this, it is possible to show that the SDE in Eq. (2) is more accurately presented as

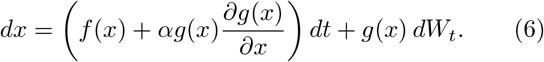

The presence of multiplicative noise can then be seen to introduce an additional term, 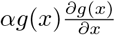, known as the *noise-induced drift* term [12]. The dependence of Eq. (6) on the stochastic integration convention *α* is also reflected in the steady-state probability distribution *P*_*s*_(*x*) of the state of the system *x*, and consequently, the probabilistic landscape *U*_*q*_(*x*): explicitly, it is known that

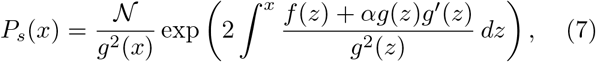

where 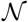 is the normalisation constant. The probabilistic landscape as computed by Eq. (4) is then given by

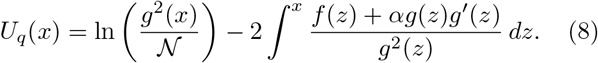

While both the Itô and Stratonovich interpretations are legitimate, Eq.’s (7) and (8) show that different choices of *α* can lead to different solutions for *P*_*s*_(*x*), and subsequently, *U*_*q*_(*x*). Thus, from a modelling perspective, an SDE is somewhat meaningless until it is assigned a stochastic integration convention. Deciding which interpretation should be adopted for a given SDE, known as the Itô-Stratonovich dilemma, has been widely debated and remains a source of controversy even today [71]. To complicate matters even further, it is possible that the stochastic integration convention required to correctly interpret an SDE may vary as the parameters of the system change [72].

In Box 2 we provide guidelines on choosing a stochastic integration convention in biological modelling. We mention that there also exist a handful of approaches that attempt to circumvent the Itô-Stratonovich dilemma. In [33], for example, the dependence on *α* is eliminated by transforming a one-dimensional Langevin equation with a multiplicative noise term to one with a constant diffusion term. Alternatively, one may wish to consider the A-type integral that extends beyond the *α*-type interpretation we discuss above, and enables a general consistency between the deterministic and stochastic dynamics. This may be useful in modelling situations where it is necessary to preserve the deterministic dynamical information in the stochastic counterpart [73].

#### Box 2 The Itô-Stratonovich Dilemma

##### The source of controversy

Stochastic differential equations (SDEs) are frequently used to model systems subject to random fluctuations. When the noise component is state-dependent (multiplicative), the resulting SDE (see Eq.(2)) is no longer well-defined without additional information: the precise behaviour depends on the adopted convention for stochastic integration. Essentially, the difficulties arise in solving the SDE and attempting to evaluate stochastic integrals of the form 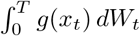. In the standard Riemann integral of integral calculus, we consider increasingly finer partitions 0 = *t*_0_ *< t*_1_ < ... < *t*_*n*_ = *T* of the interval [0, *T*] over which we wish to integrate. For each sub-interval [*t*_*k*−1_, *t*_*k*_] in a partition of [0, *T*], the integral is approximated by a rectangle of height given by a value of the integrand *g*(*x, t*) at some point 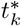 in [*t*_*k*−1_, *t*_*k*_]. The overall integral is then approximated by the sum of these rectangles – the so-called Riemann sum. Conveniently, for a smooth function, the limit over increasingly finer partitions does not depend on the precise partitions (provided the subinterval lengths go to 0) nor on the precise evaluation of *g*(*x, t*) within the intervals, so that the integral itself can be defined as the limit. For stochastic processes this is no longer true: different choices of the evaluation point within the sub-intervals can give a different outcome. Mathematically, we can express this dependence as 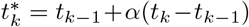, where the parameter *α* is from [0, 1] and is known as the *stochastic integration convention*. Common choices include *α* = 0 or *α* = 0.5, known as the Itô or Stratonovich interpretation, respectively.

##### Common stochastic calculus prescriptions

Rewriting an SDE, in conjunction with a stochastic convention *α*, gives Eq. (6) – and the value of *α* is at the heart of the Itô-Stratonovich debate. The Itô interpretation evaluates the integrand for the *k*^th^ interval [*t*_*k*−1_, *t*_*k*_] at *t*_*k*−1_. Thus the value at *t*_*k*−1_ is chosen based only on information available at that time; referred to as *non-anticipating*, and means that the Itô integral is a *martingale*; a useful mathematical property [11]. A consequence is that there is no correlation between *g*(*x*_*t*_) and *dWt*. In contrast, the Stratonovich interpretation evaluates the integral at 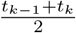, meaning that the value taken depends on the future time *t*_*k*_; it is said to be *anticipating*. This results in correlation between *g*(*x*_*t*_) and *dWt*.

##### Mathematical equivalence

For a fixed SDE, the Itô and Stratonovich interpretations yield different results. However, once an *initial* prescription has been assigned, one can easily convert between the two interpretations, either by adding or subtracting the noise-induced drift term [11, 74] (see main text). Thus, for every Itô-SDE there is a Stratonovich-SDE with identical solution. If we want to predict the behaviour of a stochastic model, we need to decide which stochastic calculus is appropriate, given they may offer different answers. Conversely, if we have observed behaviour and wish to infer the underlying dynamics of the system, then the conundrum again arises: depending on the calculus different equations are required to achieve the given solution.

##### Itô or Stratonovich in biological modelling?

The appropriate stochastic calculus to prescribe is usually chosen on physical grounds and rests upon the characteristics of the noise. Thus, when the physical properties of the stochastic fluctuations are not understood, the choice of Itô or Stratonovich becomes less clear.

*Intrinsic noise* – For chemical reaction systems, such as the Schlögl model considered in the main text, the CME is the most accurate framework for describing the full evolution of the system, including the inherent stochasticity. Upon expansion, the CME naturally leads to the well-defined CFPE, which is equivalent to the CLE; an Itô SDE [59, 75]. Thus, under the CME formalism, the choice of Itô or Stratonovich is effectively redundant [74].

*Extrinsic noise* – When modelling systems subject to extrinsic fluctuations, it is important to understand the physical properties of the noise, specifically the correlation time. Informally, the correlation time is a measure of the “memory” of the stochastic process. White noise has zero correlation time, and thus can be thought of as a completely unpredictable stochastic process. On the other hand, non-white or *coloured* noise has finite correlation time. If the extrinsic noise has *small*, but finite correlation time then a common approach is to model the noise as “approximately white”. Here the Stratonovich prescription should be adopted [76, 74]. This approach is ideal for systems that operate on time scales that are much greater than the correlation time of the noise. In some settings, however, extrinsic noise may admit non-negligible autocorrelation time. For example, in gene expression, it has been shown experimentally that some extrinsic noises have autocorrelation time comparable to the cell cycle [77, 78]. In such cases, one may wish to explicitly model the coloured extrinsic noise, by replacing the white noise term *W*_*t*_ in Eq. (2) of the main text, with some coloured stochastic process. In doing so, the SDE becomes non-Markovian and the associated FPE is not generally obtainable, although some recent progress has been made analytically with respect to additive coloured noise [79]. Nevertheless, there exist alternative approaches for simulating systems driven by coloured extrinsic fluctuations. [63, 80, 81]

A second subtlety arises in the computation of the probabilistic landscape, *U*_*q*_(*x*). In the case of additive noise, it is clear from Eq. (8), that there is no contribution from the noise-induced drift term, and so the deterministic potential will be recovered as *U*_*q*_(*x*), up to a scale and shift; this is implicit in the original justification for computing the landscape as *U*_*q*_(*x*) [16]; see also [14, 82]. In the case of multiplicative noise, however, even for a fixed stochastic integration convention *α*, the closeness of the probabilistic landscape *U*_*q*_(*x*) to the deterministic landscape *U*(*x*), will strongly and non-trivially depend on the interaction between the deterministic and stochastic dynamics. In the following section, we use a set of simple illustrative examples to demonstrate the dependence of *U*_q_(*x*) on the choice of the stochastic integration convention *α*, as well as the functions *f*(*x*) and *g*(*x*). In particular, we demonstrate and prove mathematically that for common choices of *α*, the probabilistic landscape of a bistable system (consisting of two stable fixed points) may in fact become monostable (consisting of a single fixed point) in the presence of multiplicative noise. Thus, unlike in the case where the noise component *g*(*x*) is constant, the probabilistic landscape will not necessarily reflect the deterministic potential landscape.

A further important consideration is the identifiability of the underlying deterministic dynamics in the presence of stochasticity. In the context of Eq. (7), this corresponds to asking whether or not *f*(*x*) is uniquely determined by *U*_*q*_(*x*). We provide an example that shows that there are systems characterised by strikingly different *f*(*x*) and different *g*(*x*) that give rise to the same steady-state probability distribution, and by extension, the same probabilistic landscape. Thus, it is not possible to identify the deterministic dynamics on the basis of the observed landscape *U*_*q*_(*x*) (or single-cell snapshot data) alone. Despite this, we show that for one-dimensional stochastic systems, any one of *g*(*x*), *f*(*x*) and *P*_*s*_(*x*) can be uniquely determined from knowledge of the other two. Consequently, the deterministic dynamics *f*(*x*) (equivalently, *U*(*x*)), can be determined from *U*_*q*_(*x*), only once we obtain additional information about the noise component *g*(*x*).

### Effect of noise form and intensity on the landscape

#### A double-well potential model

The model we consider here is not intended to represent any explicit biological system, but rather serves as an illustration [83] for studying the impact of noise on the stationary solutions of stochastic differential equations, and the corresponding probabilistic landscapes. While the model may be simple, it is able to give rise to intricate and biologically relevant behaviour such as bifurcations; these are an important hallmark of cell-fate decision making and mark the qualitative changes in cellular phenotype [2]. Examining how noise changes the topology^3^ of the landscape, specifically in the presence of such bifurcation points and lineage branching events, then ultimately provides insights into how it may influence cell-fate decision making. Throughout, we consider the classic case of a single particle moving in a double-well potential, and examine the effects of both a state-independent (additive) and state-dependent (multiplicative) noise component. We define the system in terms of the potential function *U*(*x*) described by,

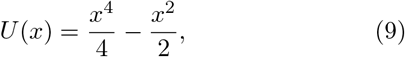

where *x* represents the state of the system i.e. the entire GRN that shapes the respective landscape. Moving *x* in state space represents a change in the gene expression profile of a cell. This one-dimensional potential is plotted in Fig. 1(B) and is reminiscent of the classic Waddington landscape, displayed in Fig. 1(A). Here the valleys that represent distinct cell states are depicted by the stable stationary points of the system, occurring at *x*_min_ = ±1, while the hill between them is depicted by the unstable stationary point at *x*_max_ = 0, and represents a transition or intermediate cell state [26, 84].

**Figure 1:**
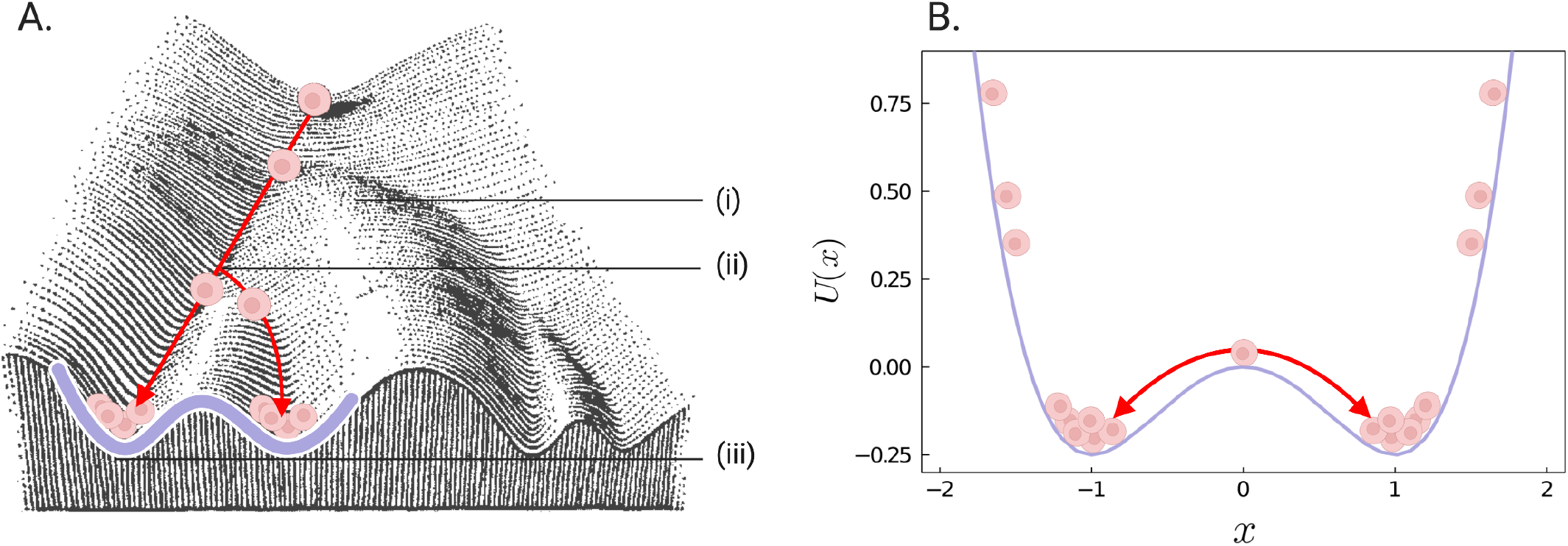
(A) Waddington’s classical representation of the epigenetic landscape where cells are described as marbles rolling downhill. The landscape is shaped by the underlying gene regulatory network. Here, we overlay cells navigating the landscape with less differentiated cells starting at the top and traversing downhill. Occasionally, a cell meets a branching point (ii) where it must decide which path to follow. This bifurcation represents a cell-fate decision point. Eventually, differentiated cells come to rest in the valleys of the landscape (outlined in purple) representing areas of low potential and high probability. The system represents a supercritical pitchfork bifurcation and is monostable (i) (stable less differentiated cell state) until it reaches a bifurcation point (ii), where after it becomes bistable (iii) (two distinct stable cell states). (B) Cells moving in the potential function *U*(*x*), representing a one-dimensional version of the landscape in (A). The figure in (A) is adapted from [3].

#### Additive noise

We consider first the case of additive noise. Here *g*(*x*) = *σ*, with noise intensity *σ >* 0. To build intuition, we begin by considering informally the effects of this noise on the potential landscape defined in Eq. (9). In Fig. 2, we plot both the potential function *U*(*x*) and the noise function *g*(*x*) = 1; these are shown as the purple and red curves respectively. We indicate with blue arrows the *net propensity* of the system’s trajectory, which we define as the net effect of noise on the potential landscape. The most probable regions of the system’s trajectory are indicated by the pink cells. As the noise is uniform across the deterministic landscape *U*(*x*), the net propensity of the system is proportional to the derivative of the potential landscape *U*(*x*). In particular, the net propensity is zero at the stationary points of *U*(*x*), and so these will be unaltered by the effects of noise. An increase in noise intensity has only a disorganising effect on the landscape i.e., it merely jostles the cells around in the potential, thereby increasing the probability of a cell to hop from one stable state to another. We now consider the system more formally.

**Figure 2:**
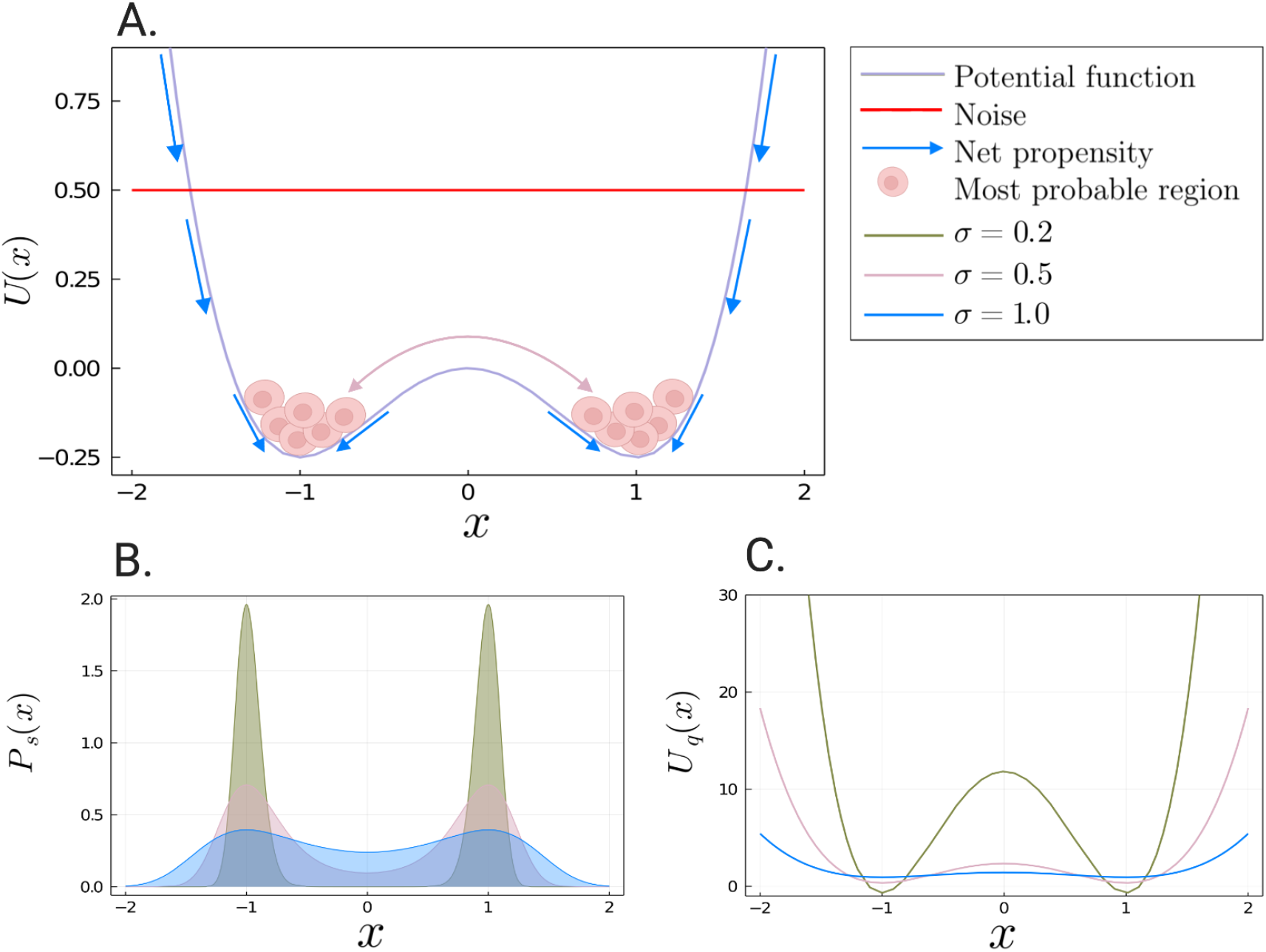
Effect of additive noise on the landscape. (A) A schematic of the system’s trajectory. We consider a deterministic double-well potential (purple curve) driven by additive noise (red curve). The blue arrows represent the net propensity of the system, with cells gathering in the most probable regions. The pink arrow indicates rare transitions from one stable state of the system to the other. (B) Analytical steady-state probability distribution *P*_*s*_(*x*) given by Eq. (10). (C) The corresponding probabilistic landscape *U*_*q*_(*x*) given by Eq. (11). Both figures are plotted for the model defined in Eq. (9) with increasing strengths of additive noise. The different colours represent the different levels of noise: the green curve is for *σ* = 0.2, the pink curve for *σ* = 0.5 and the blue curve for *σ* = 1.0. As *σ* increases both *P*_*s*_(*x*) and *U*_*q*_(*x*) flatten, making rare transitions between states more probable. The maxima of *P*_*s*_(*x*) and minima of *U*_*q*_(*x*) are preserved at *x* = *±*1.

As discussed above, both the Itô and Stratonovich interpretations of the stochastic system lead to the same stationary probability distribution. Thus, the estimated landscape *U*_*q*_(*x*) will not depend on *α* when *g*(*x*) is constant.

As *f*(*x*) is related to *U*(*x*) by way of Eq. (3), it follows from Eq.’s (7) and (8), that the stationary probability distribution, and subsequently the probabilistic landscape *U*_*q*_(*x*), are given by

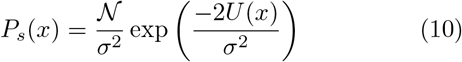

and

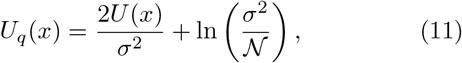

respectively. Here 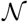 is the normalisation constant. We display the steady-state probability distribution *P*_*s*_(*x*) in Fig. 2(A), and the corresponding probabilistic landscape *U*_*q*_(*x*) in Fig. 2(B). In each figure, the three curves are computed for increasing values of noise *σ*. We see that as *σ* increases, both *P*_*s*_(*x*) and *U*_*q*_(*x*) flatten, but the maxima of *P*_*s*_(*x*) and therefore minima of *U*_*q*_(*x*) remain fixed at *x*_max_ = ±1 and *x*_min_ = ±1, respectively. Consequently, the overall topology of the landscape is preserved in the case of additive noise. That is to say the stable stationary points of the probabilistic landscape *U*_*q*_(*x*) directly correspond to those of the deterministic landscape or potential, *U*(*x*). Note that this does not depend on the specific choices of *f*(*x*) and *g*(*x*) that we have made here; *U*(*x*) will always be recovered as *U*_*q*_(*x*), albeit scaled and shifted by the noise parameter *σ*.

#### Multiplicative noise

For multiplicative noise the probabilistic landscape *U*_*q*_(*x*) is profoundly shaped by the interaction between the deterministic dynamics *f*(*x*) (or equivalently *U*(*x*)) and the stochastic dynamics *g*(*x*). We begin by considering the effects of different noise forms on our model, showing that the topological features of the deterministic landscape can be distorted by multiplicative noise. Following this, we provide a general result, which shows the fixed points of the deterministic landscape *U*(*x*) will always be preserved by *U*_*q*_(*x*), when the noise is related to the deterministic landscape by way of a power relationship. A particularly natural way in which noise might arise is when the noise component *g*(*x*) is proportional to the deterministic landscape itself. This corresponds to a special case of our general result, and we discuss how the effects of this noise on the landscape are then qualitatively indistinguishable from that of additive noise. Our result holds for *any* potential function *U*(*x*); it is not limited to any specific form such as the double-well potential considered here.

#### Example 1: Shifting the fixed points

We begin by considering the effect of the noise component 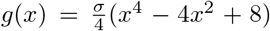, for *σ >* 0, on the model *U*(*x*) defined in Eq. (9). In order to build intuition, we display in Fig. 3(A) both the potential function *U*(*x*) and noise function *g*(*x*) (for *σ* = 1); these are the purple and red curves, respectively. We see that the noise function broadly follows the shape of the potential function, however the stable stationary points of *g*(*x*) do not agree with those of *U*(*x*): for *g*(*x*) these occur at 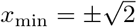, while for *U*(*x*) these occur at ±1. The net propensity of the system is indicated by blue arrows, and the most probable regions of the system’s trajectory are represented by the pink cells. As the noise is much higher at the unstable stationary point (*x*_*min*_ = 0) relative to the strength of noise to the left and right of it, cells are pushed away from the unstable point. The net propensity of the system is therefore non-negligible at the stable stationary points of *U*(*x*), and so the system’s trajectory is more often found in the region slightly beyond the stable stationary points of *U*(*x*). The resulting stationary distribution is therefore bimodal, with modes occurring beyond *x*_max_ = ±1. We now formalise this intuition.

**Figure 3:**
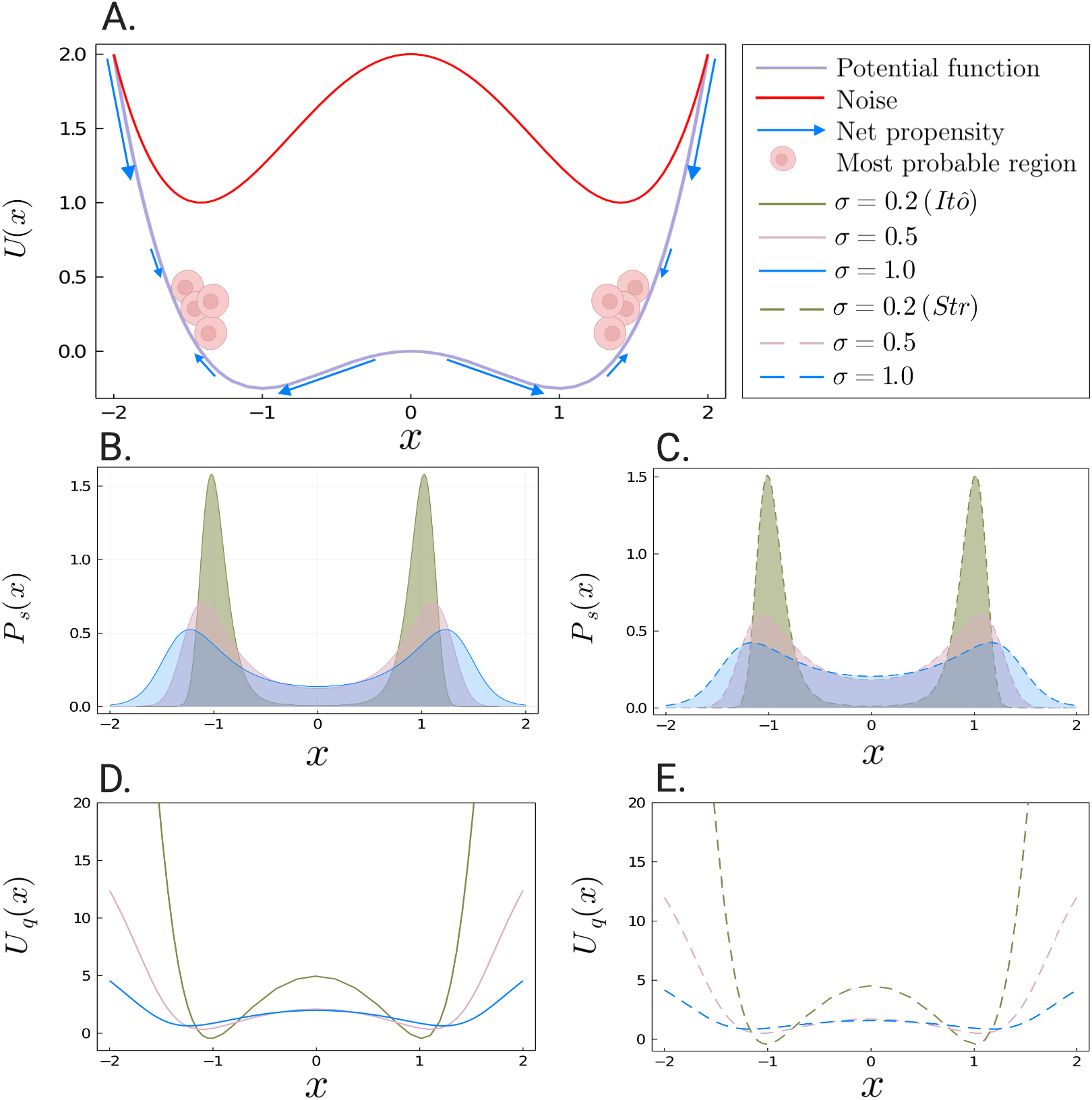
Effect of multiplicative noise on the landscape: shifting fixed points. (A) A schematic of the system’s trajectory. Here we consider a deterministic double-well potential driven by multiplicative noise. The net propensity of the system (as represented by the blue arrows) is to drive cells *away* from the origin. Cells are then more often found beyond the stationary points of the deterministic landscape *U*(*x*). (B&C) Analytical steady-state probability distribution *P*_*s*_(*x*) given by Eq. (12), for *α* = 0 (solid lines) and *α* = 0.5 (dashed lines), respectively. (D&E) The corresponding landscape *U*_*q*_(*x*) given by Eq. (13), for *α* = 0 (solid lines) and *α* = 0.5 (dashed lines), respectively. Figures (B-E) are plotted for the model defined in Eq.(9) with increasing strengths of multiplicative noise. Here *σ* = 0.2 is represented as the green curve, *σ* = 0.5 as the pink curve, and *σ* = 1.0 as the blue curve. As *σ* increases, the maxima of *P*_*s*_(*x*) and (resp. minima of *U*_*q*_(*x*)) are shifted away from the origin towards the fixed points of the noise curve *g*(*x*). As *σ* increases both *P*_*s*_(*x*) and *U*_*q*_(*x*) flatten.

As the noise function *g*(*x*) is state-dependant, the stochastic system may be interpreted in either the Itô or Stratonovich sense, leading to different solutions for the steady-state probability distribution *P*_*s*_(*x*) and, subsequently, the probabilistic landscape *U*_*q*_(*x*). The dependence on the stochastic integration convention *α* can be seen from Eq.’s (7) and (8) with *f*(*x*) = *x* − *x*^3^ and 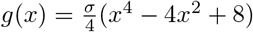 explicitly we have

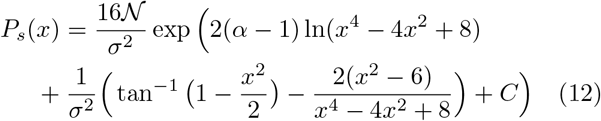

and

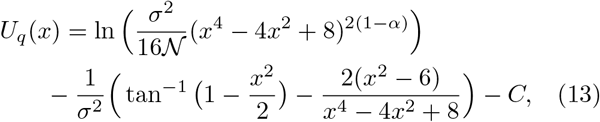

where 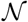 is the normalisation constant and *C* is the constant of integration.

In Fig. 3, we plot the steady-state probability distribution *P*_*s*_(*x*) and the corresponding landscape *U*_*q*_(*x*), for both of the Itô (*α* = 0) and Stratonovich (*α* = 0.5) interpretations. We use solid lines to represent the Itô interpretation (Fig. 3(B&D)), while dashed lines represent the Stratonovich interpretation (Fig. 3(C&E)). Each figure is plotted for increasing strengths of multiplicative noise. As with additive noise, an increase in *σ* broadens the steady-state probability distribution *P*_*s*_(*x*), leading to a decrease in the depth of the wells of the corresponding landscape *U*_*q*_(*x*). The modes of the distribution (resp. wells of the landscape) however do not remain fixed: they are now shifted away from the origin. This displacement in the modes (wells) occurs for both of the stochastic interpretations. In the limit of large noise (i.e. *σ* → ∞), the modes of *P*_*s*_(*x*) (wells of *U*_*q*_(*x*)) approach the stable stationary points of the noise function *g*(*x*), which occur at 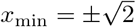.

While the effects of the noise *g*(*x*) on the deterministic potential are visually almost indistinguishable for the Itô (Fig. 3(B&D)) and Stratonovich (Fig. 3(C&E))) interpretations of the stochastic system, under close inspection it can be seen that the Stratonovich interpretation has produced slightly flatter curves. In fact, they are also slightly narrower. For any given noise intensity *σ*, the modes of *P*_*s*_(*x*) (resp. wells of *U*_*q*_(*x*)) are shifted further away from the origin under the Itô interpretation, in comparison to the Stratonovich interpretation. The broadening of *P*_*s*_(*x*) (resp. flattening of *U*_*q*_(*x*)) however, is more pronounced under the Stratonovich interpretation, for any given noise intensity *σ*.

A similar example can be constructed using the noise component 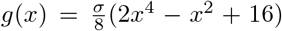. Again, this noise curve broadly follows the shape of the potential landscape *U*(*x*), but has minima occurring at 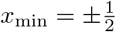. This noise now shifts the stationary points of the landscape *towards* the unstable stationary point at the origin, which approach 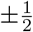 in the limit of large noise. Details of this example can be found in the supplementary material.

#### Example 2: Creation and destruction of fixed points

We next consider the effect of the noise component *g*(*x*) = *σ*(1 + *x*^2^), for *σ >* 0 on the double-well potential *U*(*x*). We begin by considering informally the effects of this noise on the system. In Fig. 4(A), we plot both *U*(*x*) and the noise function *g*(*x*) as the purple and red curves, respectively. As the noise is much higher at the left- and right-most extremes of the landscape, relative to the strength of noise at the origin, cells are pushed towards the unstable stationary point of *U*(*x*). The net propensity of the system (indicated by the blue arrows) is non-negligible at the stable stationary points of *U*(*x*) (*x*_*min*_ = ±1), and so the system’s trajectory is more often found either side of the unstable point, now closer to the origin. For lower noise intensities, the resulting steadystate probability distribution will therefore be bimodal, with modes occurring between (−1, 1). When the noise intensity becomes large enough, eventually the two modes (corresponding to the most probable regions of cells) will collide at the origin, resulting in a unimodal distribution.

**Figure 4:**
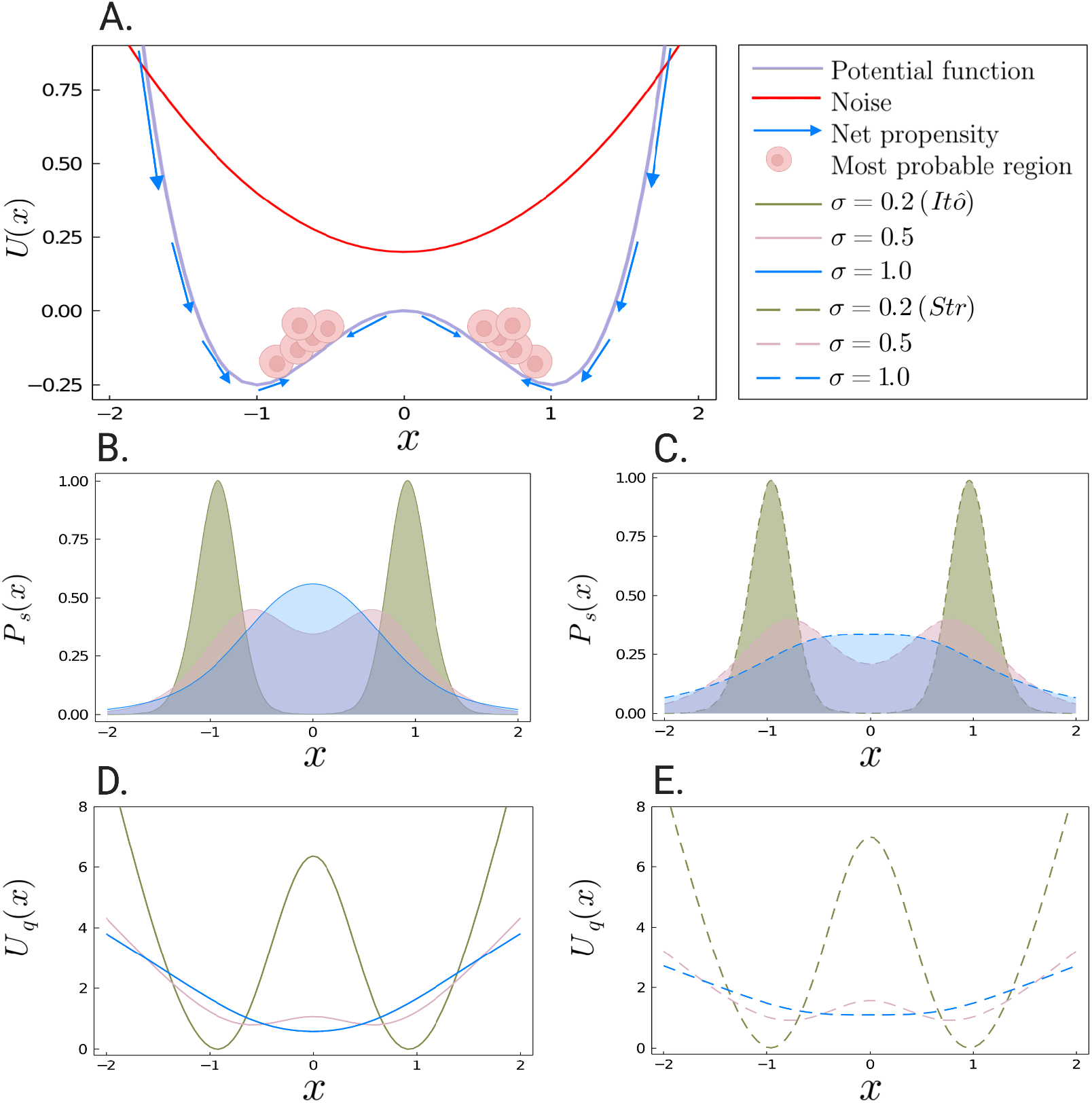
Effect of multiplicative noise on the landscape: destruction of fixed points. (A) A schematic of the system’s trajectory. We consider a deterministic double-well potential driven by multiplicative noise. The net propensity (blue arrows) pushes cells *toward* the origin. (B&C) Analytical steady-state probability distribution *P*_*s*_(*x*) given by Eq. (14), for *α* = 0 (solid lines) and *α* = 0.5 (dashed lines), respectively. (D&E) The corresponding landscape *U*_*q*_(*x*) given by Eq. (15), for *α* = 0 (solid lines) and *α* = 0.5 (dashed lines), respectively. Figures (B-C) are plotted for the model defined in Eq. (9) with increasing strengths of multiplicative noise. Here *σ* = 0.2 is represented by the green curve, *σ* = 0.5 by the pink curve, and *σ* = 1.0 by the blue curve. As *σ* increases, the maxima of *P*_*s*_(*x*) and minima of *U*_*q*_(*x*) are shifted towards the origin until eventually they collide (blue curve). As *σ* increases both *P*_*s*_(*x*) and *U*_*q*_(*x*) flatten.

Again, as the noise is state-dependant, the stochastic system may be interpreted in either the Itô or Stratonovich sense. Thus, the steady-state distibution *P*_*s*_(*x*) and, therefore, the probabilistic landscape *U*_*q*_(*x*) will depend on *α*. To see this explicitly, we consider Eq.’s (7) and (8) with *f*(*x*) = *x* − *x*^3^ and *g*(*x*) = *σ*(1 + *x*^2^); we have

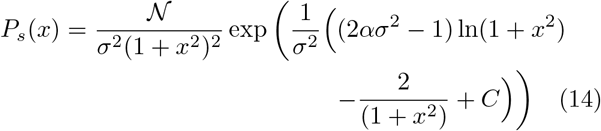

and

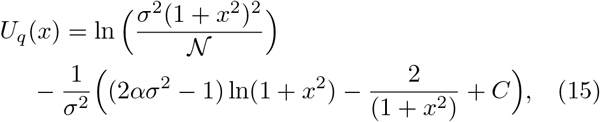

where 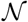 is the normalisation constant and *C* is the constant of integration.

In Fig. 4 we plot the steady-state probability distribution *P*_*s*_(*x*) and the corresponding landscape *U*_*q*_(*x*), for both stochastic integration conventions *α* = 0 (Fig. 4(B&D) and *α* = 0.5 ((Fig. 4(C&E)). Again, each figure is plotted for the model in Eq. (9), with increasing strengths of multiplicative noise. Solid lines represent the Itô interpretation (*α* = 0), and dashed lines represent the Stratonovich interpretation (*α* = 0.5). As the noise intensity *σ* increases, we see a broadening in the steady-state probability distribution *P*_*s*_(*x*), leading to a decrease in the depth of the wells of the corresponding landscape *U*_*q*_(*x*). The shape of the probabilistic landscape *U*_*q*_(*x*), however has a major departure from that of the potential *U*(*x*). More specifically, the modes of the steady-state probability distribution, and subsequently the minima of the probabilistic landscape are displaced towards the origin, as the noise intensity *σ* increases. The extent of these effects vary according to stochastic integration convention (*α* = 0 or *α* = 0.5). This can be seen explicitly from the minima of the probabilistic landscape *U*_*q*_(*x*), which are given by

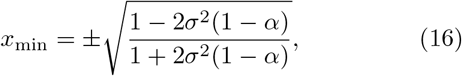

Unlike in the case of additive noise, the minima of *U*_*q*_(*x*) agree with those of the potential landscape *U*(*x*) only in the limit as *σ* approaches 0 (i.e. the weak noise limit). While for 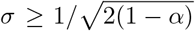, the stable stationary points (*x*_min_) of *U*_*q*_(*x*) collide, resulting in a single stable stationary point at the origin. We refer to the point

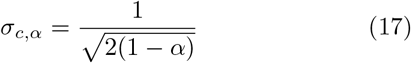

at which the system transitions from bistable to monostable as the *critical noise*. Notably, this point depends on the stochastic interpretation *α*. For the Itô interpretation, the critical noise 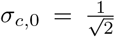, while for the Stratonovich interpretation *σ*_*c*,0.5_ = 1. Thus, under the Stratonovich interpretation, a higher noise intensity is required to drive the system from bistable to monostable. The discrepancy between the stochastic interpretations can also be seen in Fig. 5, where *U*_*q*_(*x*) is plotted for both interpretations with noise intensity 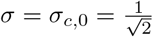.

In this example, we have demonstrated that the probabilistic landscape of a bistable system may in fact become monostable in the presence of multiplicative noise. In other words, multiplicative noise can destroy the fixed points of a system. Later, we will see how multiplicative noise can also drive the creation of fixed points: a monostable system under the influence of multiplicative noise can become bistable. Such phenomenon are known as *noise-induced transitions* and have been demonstrated in a number of biological systems, including gene expression [85, 86, 87], enzymatic cycles [88], cell cycle regulation [89], *E. Coli* [90], as well as synthetic biological circuits such as the toggle switch [91]. It is precisely these observations that have important consequences for dynamical systems undergoing a bifurcation in the presence of multiplicative noise. Specifically, the stochastic system *U*_*q*_(*x*) may display qualitatively different behaviour from the deterministic system *U*(*x*). We discuss bifurcations of stochastic dynamical systems in more detail below.

**Figure 5:**
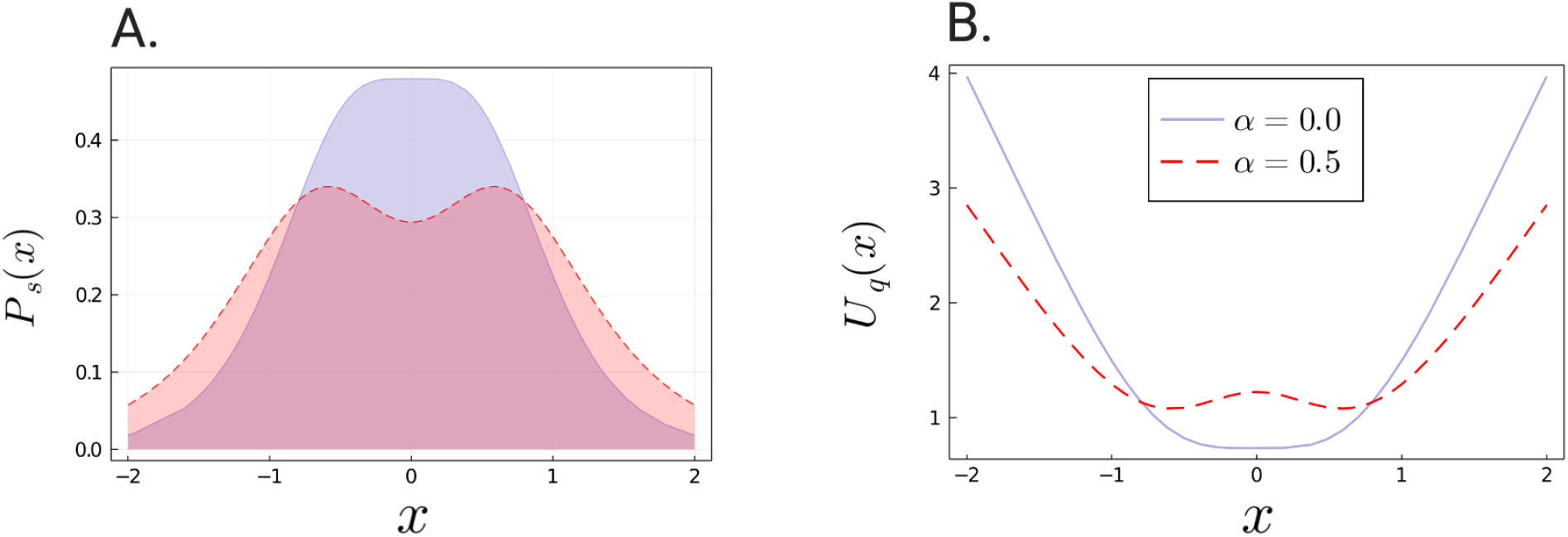
Discrepancy in the steady-state probability distribution (and corresponding landscape) for different stochastic conventions. (A) Analytical steady-state probability distribution *P*_*s*_(*x*) given by Eq.(14) and (B) the corresponding landscape *U*_*q*_(*x*) given by Eq.(15), plotted for a fixed noise intensity *σ* = 0.7. The solid line is in the Itô interpretation and the dashed line in the Stratonovich interpretation. The displacement of the maxima of *P*_*s*_(*x*), and subsequently the minima of *U*_*q*_(*x*), depend on the prescribed calculus. At *σ* = 0.7 the Itô interpretation is monostable while the Stratonovich interpretation is still bistable. Both prescriptions give rise to a drift when compared with the deterministic dynamics, although the drifts are weighted differently.

### Effect of coupled noise on the landscape

So far we have considered noise forms that are uncoupled to the underlying deterministic dynamics i.e. the noise component *g*(*x*) is independent of the deterministic component *f*(*x*). In a biological context, this approach is justified for systems with high copy-numbers of all the constituent species. For such systems, intrinsic fluctuations may be ignored and the system dynamics can be modelled as a set of deterministic rate equations. Many biological systems, however, involve molecular species with low copy-numbers, and therefore the internal randomness in the dynamics may have a significant impact on the properties of the system. The default approach for modelling intrinsic noise is by way of the CLE [59], which imposes a dependence on the drift and diffusion terms. A natural question then arises: do the results of the above analysis rely upon the independence of the deterministic and stochastic dynamics? To investigate this question, we consider the effect of fluctuations arising only from intrinsic noise on the landscape. We demonstrate our results using a simple chemical reaction network model, the Schlögl model. In our analysis, the noise component represents intrinsic noise only, and is coupled to the underlying deterministic dynamics. We demonstrate here that this constraint on the form of the noise component *g*(*x*) can also have significant impacts on the properties of the system, and hence the quasi-potential landscape.

The Schlögl model, originally introduced in 1972, is the canonical example of a chemical reaction system that exhibits bistability [92]. The reactions of this model are

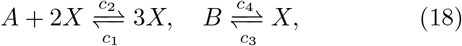

where *A, B* are chemical species with constant concentrations *a, b*, respectively, and *x* is the concentration of the dynamic chemical species *X*. Assuming mass-action kinetics, the rate equation describing the deterministic dynamics of this system is given by,

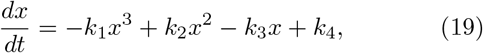

where for notational convenience we let *k*_1_ ≔ *c*_2_, *k*_2_ ≔ (*a*+3)*c*_1_, *k*_3_ ≔ *ac*_1_ +2*c*_2_ +*c*_4_, and *k*_4_ ≔ *bc*_3_. Depending on the parameters, the deterministic system will exhibit one or two stable states, as determined by the discriminant of the cubic polynomial *f*(*x*) = −*k*_1_*x*^3^ +*k*_2_*x*^2^−*k*_3_*x*+*k*_4_; for a more detailed picture of the deterministic kinetics see [93, 94]. The potential function, as determined by Eq. (19), is given by,

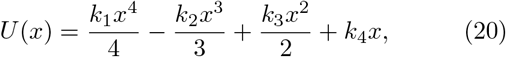

In Figure 6(A) (left plot), we display the one-dimensional potential function *U*(*x*) for the Schlögl reaction network in the unistable case; this is represented by the purple curve. The system has exactly one stationary stable point, which occurs at *x*_min_ ≈ 8.63. We note that while visually there appears to be a second minimum in the potential, mathematically, this is not a stationary point.

**Figure 6:**
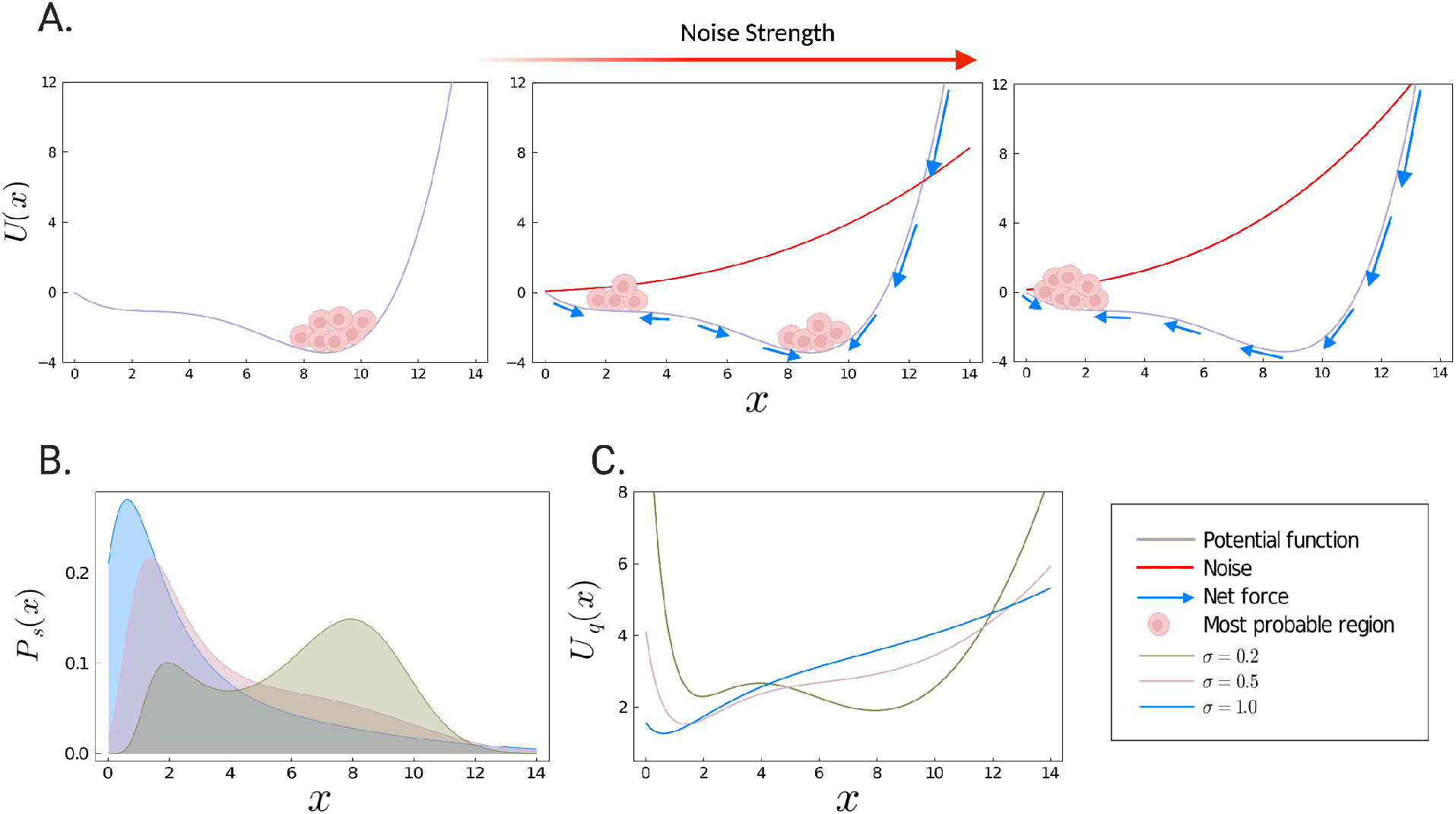
Effect of intrinsic noise on the Schlögl model. (A) A schematic of the system’s trajectory. We consider a bistable deterministic potential for the Schlögl reaction network driven by intrinsic (Langevin) noise. The net propensity (blue arrows) pushes cells *toward* the origin, and away from the right-hand stable point. (B) Analytical steady-state probability distribution *P*_*s*_(*x*) given by Eq. 22. (C) The corresponding landscape *U*_*q*_(*x*) as computed by Eq. (8). Figures (B-C) are plotted for the Schlögl model with increasing strengths of noise. Here *σ* = 0.2 is represented by the green curve, *σ* = 0.5 by the pink curve, and *σ* = 1.0 by the blue curve. The steady-state distribution and *P*_*s*_(*x*) transitions from unistable (no noise) to bimodal (the green curve), and back to unimodal (pink and blue curves) again, with increasing noise parameter *σ*. The effect on the landscape is thus similar: we see a transition from unistable to bistable, and then back to unistable again.

In the case of the Schlögl model, the CLE has been shown to be an accurate method for capturing temporal fluctuations that are due to the inherent stochasticity of the chemical process itself (intrinsic noise) [93]. One possible representation of the CLE for the Schlögl model is given by

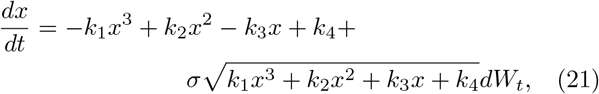

where *dW*_*t*_ is an increment in the Wiener process. Note that this representation involving only one Wiener process is equivalent to the standard CLE representation [95]. Here we include an additional noise parameter *σ* that indicates the strength of the intrinsic noise, corresponding to the level of burstiness in the system. Observe that the noise component 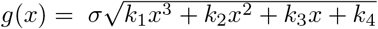 is dependent upon the deterministic component *f*(*x*). We begin our analysis of the Schlögl reaction network by first considering informally the effects of this coupled noise on the deterministic system. In Fig. 6(A), we represent the deterministic potential *U*(*x*) and the noise function *g*(*x*) as the purple and red curves, respectively. In the absence of intrinsic noise in the system (left plot), we can see that in steady-state conditions, all cells sit near to the minimum in the potential. For low noise intensities (middle plot), the net propensity is large enough to push cells into the shallow basin in the landscape, however is not large enough to overcome the deterministic force at the stable stationary point. Hence in these cases the steady-state probability distribution is bimodal. For high noise intensities (right plot), the system is more often found in a single region close to the origin. This occurs because the noise is much higher at the right-most extreme of the landscape, relative to the strength of noise at the origin, and so cells are strongly pushed into a single region close to the origin. Thus, for high noise intensities the steady-state distribution is again unimodal. Overall the system displays very interesting behaviour: deterministically the system is unistable, but with increasing levels of intrinsic noise, the system transitions to bistable, and then back to unistable again.

As the CLE is an Itô stochastic differential equation, we compute the steady-state probability distribution from Eq. (7) with *α* = 0; this gives

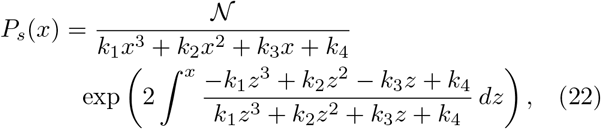

where 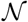 is the normalisation constant. In Fig. 6 (B&C), we plot the steady-state distribution *P*_*s*_(*x*), and the corresponding probabilistic landscape *U*_*q*_(*x*), respectively. As is expected from our informal analysis, we see a transition in the deterministic unimodal steady-state probability distribution to bimodal, and then back to unimodal, with increasing noise intensity *σ*(and thus a transition in the corresponding landscape from unistable to bistable and back to unistable again). Similar properties of the Schlögl model dynamics have been observed before [93, 94, 96].

Modelling intrinsic noise by way of the CLE is a valid approach that naturally lends itself to the Itô interpretation. Using this framework, we have shown that the Schlögl reaction model can exhibit noise-induced transitions. It is not uncommon however, for intrinsic variability to be modelled via Langevin-type equations involving the addition of ad hoc, often additive, noise terms. This approach is primarily justified on the basis of simplifying the analysis of the dynamical system, and does not follow from any physical argument (in contrast to Gillespie’s derivation of the CLE). To elucidate the limits of this approach, we refer the reader to the supplementary material for a demonstration of the effects of additive noise on the deterministic dynamics of the Schlögl model. As expected from Eq. (11) above, additive noise does not lead to qualitative changes in the landscape: the effect is only to scale and (vertically) shift the landscape, thereby preserving the position and number of fixed points. Our analysis then raises two important points: (1) intrinsic noise alone can lead to significant differences in the probabilistic and deterministic landscapes, and (2) when intrinsic noise is neglected, or is replaced by an (physically unfounded) additive noise term, the mesoscopic behaviour of the system is not reflected in the probabilistic landscape. Indeed, this extends to any arbitrary choice of a noise term in Eq. 2. The inclusion of such terms may not always be valid, and can potentially lead to spurious results in the system dynamics. The limitations we have raised here may be an important consideration for systems where it is unknown whether or not all of the species’ molecular numbers are sufficiently large enough to ignore intrinsic fluctuations.

On the other hand, many signalling systems in biology are able to accurately respond to external perturbations and robustly carry out their functions. This adaptive behaviour has been linked to the system’s ability to attenuate noise, and is often reflected in the design of GRNs [97]; feedback loops, for example, have been shown to play an important role in the system’s response to noise. While it has been shown that there are limits on a network’s ability to suppress such molecular fluctuations [98], for some networks the typical variation in the system will be small, and thus may not be amenable to the bifurcation analysis presented above. The assumption of small noise amplitude may also be reasonable when the number of molecules in the system is large. An alternative framework for capturing the behaviour of stochastic systems subject to small random perturbations is the so-called large deviation theory (LDT) of Freidlin and Wentzell [99]. The LDT provides a tool for computing the quasi-potential landscape, and for explicitly quantifying the probability of rare events, such as transitions (or switching) between metastable states. This has been employed in a number of biological contexts, including in genetic switching models [100], and yeast cell-cycle dynamics [101].

In summary, we have demonstrated the well-observed phenomenon that uncoupled (or extrinsic) multiplicative noise may have substantial effects on the dynamics of the system. In particular, we have seen that an external multiplicative noise term can destroy or create fixed points in the landscape. We have further demonstrated that coupled (or intrinsic) noise may too exhibit similar behaviour. This is to a certain extent unsurprising, considering that intrinsic noise as described by the CLE is multiplicative. To complete this section, we show that there are natural instances where multiplicative noise may in fact preserve the position and number of fixed points, in the same way that additive noise preserves these topological features of the landscape.

#### Preservation of fixed points under multiplicative noise

Let *U*(*x*) be any one-dimensional potential function, and assume that *g*(*x*) is related to *U*(*x*) by way of the following power relationship,

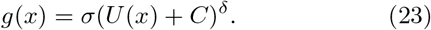

Here *σ >* 0 is the noise intensity and *C, δ* are any non-negative real numbers. Further assume that *U*(*x*)+*C >* 0, for all *x*, so that the noise component *g*(*x*) is strictly positive. Then it can be shown that, for any stochastic integration convention *α*, the fixed points of the approximated landscape *U*_*q*_(*x*) correspond precisely to those of *U*(*x*) (when they exist). Thus, for multiplicative noise of the form given in Eq. (23), the topological structure of *U*(*x*) is preserved by the probabilistic landscape *U*_*q*_(*x*), in a similar way to that of additive noise. Indeed, additive noise is a particular case of Eq (23), corresponding to when *n* = 0. The proof of this general result is provided in the supplementary material.

A natural noise form that satisfies the broader features of noise observed in cellular differentiation arises when *g*(*x*) is proportional to the deterministic landscape itself. Here, the noise component *g*(*x*) closely follows the shape of the landscape, and thus will tend to be high at regions of higher potential (unstable states), and comparatively lower at the minima in the landscape. In the context of Eq. (23) above, this corresponds to the particular case where *C* = 0 and *δ* = 1. An example demonstrating the effect of this noise on the model defined in Eq. (9) can be found in the supplementary material; the effects on the stationary distribution and probabilistic landscape can be seen to be qualitatively identical to that of additive noise. From a modelling perspective, it is challenging to justify any particular noise form, other than to ensure it meets the broader features of noise that have been experimentally observed. Our aim here is to highlight that there are more natural, and perhaps more realistic noise forms, when compared with additive noise, that are able to preserve the topology of the underlying deterministic landscape.

### Identifiability and limitations to inferring dynamics

As discussed above, a stochastic dynamical system is in general characterised by a stochastic integration convention *α*, in unison with a deterministic component *f*(*x*) and stochastic component *g*(*x*). In practise however, it is rarely the case that these components are known: experimental data is of the form *P*_*s*_(*x*), which provides only a static picture of the long-term, average dynamics of the system. Thus, some of the dynamical information of the system is lost in *P*_*s*_(*x*). How much then can we deduce of the underlying stochastic dynamics (*f*(*x*) and *g*(*x*)), as well as the stochastic calculus *α*, from the measured stationary distribution *P*_*s*_(*x*), or equivalently, the observed landscape *U*_*q*_(*x*)? This presents two possible challenges.

- The first is the possibility that there are systems characterised by different *f*(*x*) and different *g*(*x*) that yield the same stationary probability distribution *P*_*s*_(*x*), and therefore, the same *U*_*q*_(*x*).
- A second challenge is that even if the deterministic system *f*(*x*) is known, it may be possible that different choices of the noise component *g*(*x*) could give rise to the same distribution *P*_*s*_(*x*). And similarly, there may be different deterministic systems *f*(*x*) that can yield the same stationary distribution *P*_*s*_(*x*), by way of a fixed noise component *g*(*x*).

It may not be surprising that a given stationary probability distribution can represent many different dynamical systems; indeed this has been previously shown empirically [26, 40]. Nevertheless, it will be instructive to consider this more formally. Here we show that for one-dimensional stochastic systems, the steady-state distribution *P*_*s*_(*x*) can always be obtained from systems characterised by different *f*(*x*) and different *g*(*x*), for any given stochastic integration convention *α*. We demonstrate this result using an illustrative example, showing there are non-identifiability instances of the first kind (the first item above). We then discuss how non-identifiabilty can be resolved, thereby showing there do not exist non-identifiability instances of the second kind (the second item above).

#### Decomposition of the stationary probability distribution

Fix some stationary probability distribution *P*_*s*_(*x*). Then for any noise component *g*(*x*), and stochastic integration convention *α*, the deterministic dynamics *f*(*x*) can be solved as

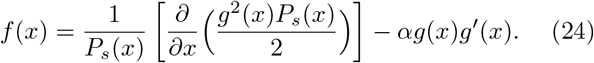

On the other hand, given any deterministic dynamics *f*(*x*), the noise component *g*(*x*) can be solved as

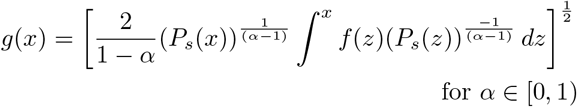

and

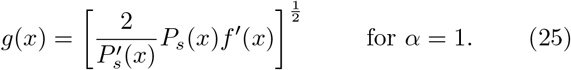

Thus, the stationary probability distribution *P*_*s*_(*x*) can be obtained by any noise component *g*(*x*) with the deterministic dynamics in Eq. (24), or by any deterministic component *f*(*x*) and the multiplicative noise given in Eq. (25); the case where *α* = 0 (the Itô interpretation) can be found in Sura et al. [102].

#### Non-identifiability of the first kind

In order to simplify our example, we will assume throughout that the stochastic integration convention *α* is known to be 0, corresponding to the Itô interpretation. In practise, the stochastic interpretation *α* is often chosen *a priori* on the basis of the available information, but in general is not known and cannot be inferred from experimental data: there are different *f*(*x*) and *α* that yield the same stationary probability distribution [11, 71, 74].

Consider the probability distribution obtained from the deterministic dynamics *f*_1_(*x*) = −*x* and the noise component *g*_1_(*x*) = *σ* exp(−*x*^2^),

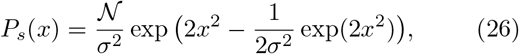

where 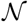 is the normalisation constant. This probability distribution is displayed in Fig. 7(A) for *σ* = 2; note that the most probable states occur at 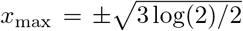 (≈ ±1.02). The corresponding potential function *U*_1_(*x*) = *x*^2^ and noise function *g*_1_(*x*) are shown in Fig. 7(B) as the purple and red curves, respectively. The net effect of noise on the potential function (as indicated by the blue arrows), as well as the most probable regions of the system’s trajectory are also shown. As the noise is much higher at the stable stationary point relative to the strength of noise to the left and right of this point, cells are pushed away from the stable point, and the system’s trajectory is more often found either side of the stable stationary point (the regions indicated by the pink cells). The result is the bimodal probability distribution given in Eq. (26).

Using the stationary probability decomposition above (Eq. (24)), we are able to obtain a second system that gives rise to the same probability distribution given in Eq. (26). This system has substantially different dynamics: the underlying deterministic dynamics 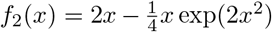 gives rise to the bistable system *U*_2_(*x*) shown in Fig. 7C; note that the stable points of *U*_2_(*x*) occur at 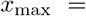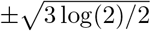, which agree with the maxima of *P*_*s*_(*x*). Moreover, the system is now perturbed only by the additive (constant) noise *g*_2_(*x*) = 1 (as opposed to the multiplicative noise *g*_1_(*x*)). In contrast to Fig. 7B, we can see that the noise acts uniformly on the system’s trajectory, and thus preserves the two stable points of the system. The result of the noise *g*_2_(*x*) on the potential *U*_2_(*x*) again produces the bimodal probability distribution given in Eq. (26).

**Figure 7:**
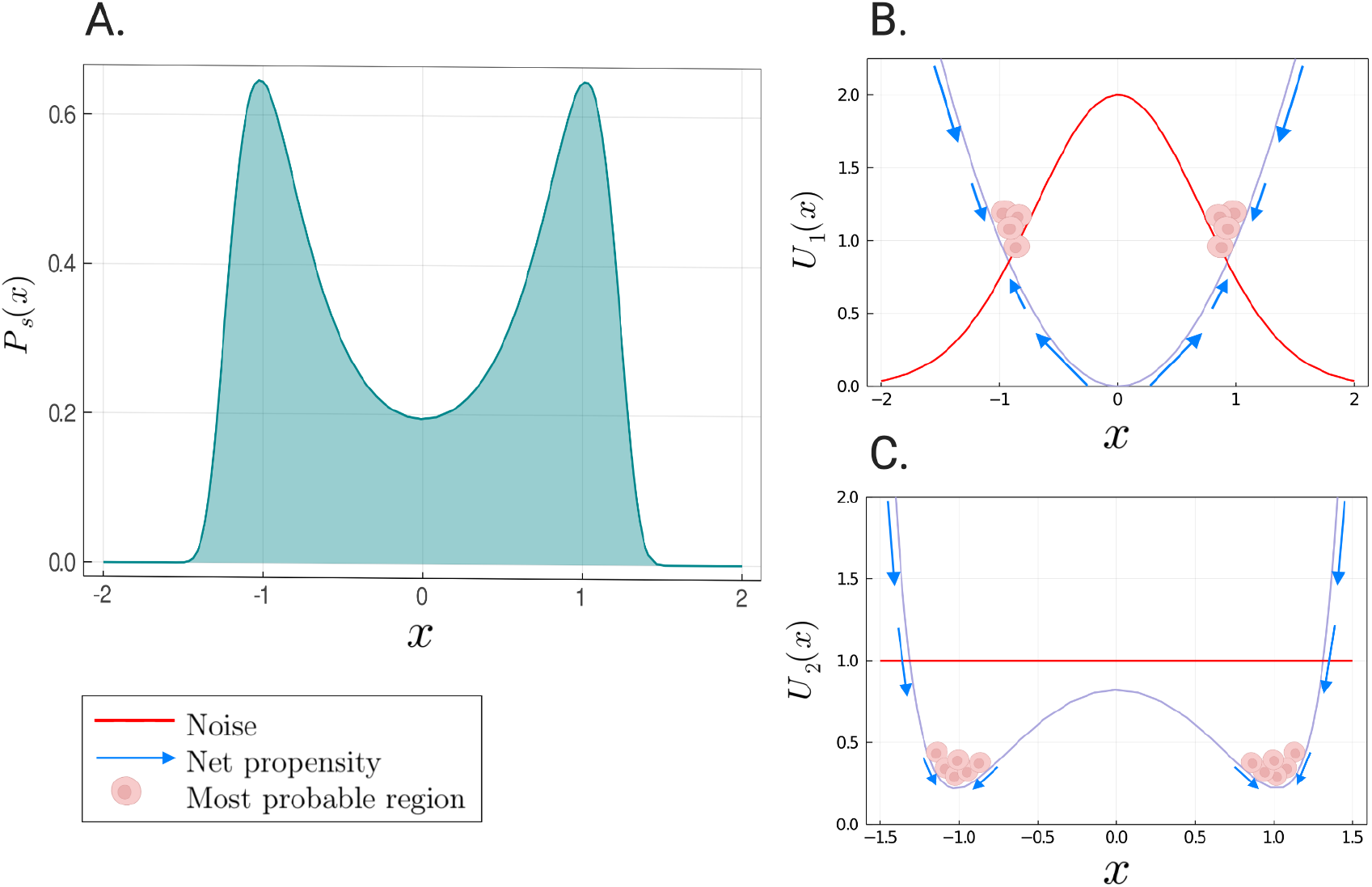
Non-identifiability of the first kind. The same bimodal probability distribution (A) arises from a unimodal system subject to multiplicative noise (B), and from a bimodal system driven by additive noise (C), resulting in non-identifiability of the first kind. In (B) the noise strength (red curve) is greater around the origin thereby pushing cells *away* from the origin and generating a bimodal probability distribution. (C) Additive noise does not shift the minima of *P*_*s*_(*x*) and so the distribution remains bimodal.

Our example shows that having knowledge of the measured stationary distribution alone (or equivalently, the observed landscape) is not enough to infer the underlying regulatory dynamics of the stochastic system. Did the stationary distribution arise from a system with only one cell state perturbed by multiplicative noise, or a bistable system with only additive noise? In biological terms, knowing the typical distribution of cell states offers very little information about *how* a cell differentiates and travels (in terms of gene activity) to its cell fate.

#### Non-identifiability of the second kind

We now consider the second identifiability problem. To simplify our consideration, we again assume the stochastic integration convention *α* is known. Given the stationary probability distribution *P*_*s*_(*x*), and *one* of the components *f*(*x*) or *g*(*x*), we are then interested in whether we can uniquely identify the remaining component using only this information. The stationary probability decomposition above tells us (almost trivially) that knowledge of one of the components is sufficient to uniquely identify the other; uniqueness follows from the uniqueness property of first-order differential equations [103] (refer to the derivations of Eq.’s (24) and (25) in the supplementary material). We now briefly demonstrate this in the context of our previous example.

Assume again that we are given the stationary probability distribution in Fig. 7 (in practise, this distribution can be obtained from experimental data). Assume further that we now have some confidence in the underlying dynamics *f*(*x*) of the system. For example, we have reason to believe the underlying system is bistable, and is described by 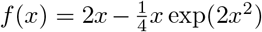. Then from Eq. (25) the noise component *g*(*x*) can be uniquely solved as

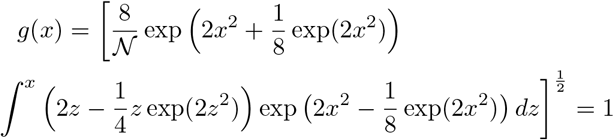

Thus, for one-dimensional stochastic systems, any one of *g*(*x*), *f*(*x*) and *P*_*s*_(*x*) can be determined from knowledge of the other two. We remark that we have considered here only gradient systems (i.e. systems satisfying the relationship in Eq. (1)).

As mentioned above, high-dimensional systems found in biology may not arise as pure gradient systems. That is, the deterministic portion of the dynamics includes an additional non-gradient component, the curl flux. Recent work has shown that this non-gradient force plays a prominent role in driving the dynamics of non-equilibrium systems, and subsequently in shaping the landscape [104, 105]. The addition of a non-gradient component to the drift introduces further challenges to the identification of the regulatory dynamics based upon (static) experimental data alone. In particular, there are now non-identifiabilty instances of the second kind. In [26] for example, it is shown that even when the noise component is fixed, there are systems defined by gradient dynamics and systems defined by non-gradient dynamics that admit qualitatively indistinguishable steady-state probability distributions. Although some progress towards the full identification of the regulatory dynamics has been made, in both the equilibrium [40] and non-equilibrium settings [42], there is still more work to do. Indeed, the aforementioned reconstruction methods are limited to potential driven systems with additive noise.

### Stochastic bifurcations

We have seen how noise may change the landscape qualitatively at critical parameter values: for example, transitioning from a bistable system to a monostable system. These changes can be viewed as binary branches on the Waddington landscape where cell-fate decisions take place, and may provide insight into how noise affects cell-fate decision making. At each branching point, a cell adopts one of two possible cell states [106], each defined by a unique gene expression profile or GRN. Mathematically, this point can be viewed as a *bifurcation* [2, 36, 107, 108]. For deterministic dynamical systems a bifurcation results in a qualitative change of the solution (nature and number of fixed points) as a parameter, or set of parameters are varied [109].

While bifurcation theory for deterministic dynamical systems is well-developed (and indeed extensively studied), the extension to its stochastic counterpart is perhaps less so. In stochastic dynamics there is potential for richer dynamics (see e.g. [110] and [111]), and we commonly distinguish between two types of bifurcations [112]: *D-bifurcations*, are the analogues of deterministic bifurcations; the ‘D’ stands for ‘dynamical’ and this type of bifurcation reflects sudden changes in the sign of the leading Lyapunov exponent of the dynamics [113]. *P-bifurcations*, by contrast, reflect sudden changes in the steady-state probability distribution over states. We have already seen examples of P-bifurcations: in the multiplicative noise examples above, we have seen how the probability density can transition from bimodal to unimodal, or even from unimodal to bimodal, and back again, only by varying the noise intensity parameter. These changes, known as noise-induced transitions, indicate that the system around the P-bifurcation point is not structurally stable [112, 6], and show how the stochastic system can display qualitatively different behaviour from that of the deterministic system.

It is possible for D- and P-bifurcations to occur individually or jointly, and the interplay between these two types of qualitative change in system behaviour is generally difficult to disentangle. Baxendale [114] for example, gives a dynamical system exhibiting a stochastic D-bifurcation, while the steady-state probability distribution is independent of the bifurcation parameter. On the other hand, Crauel and Flandoli [115] consider the stochastic system obtained from Eq. (9) in the presence of additive noise, and show that while the stationary probability density has a pitchfork bifurcation of equilibria, the dynamical structure of the system remains stable (i.e. the system does not exhibit a D-bifurcation).

As static measures are unable to capture the full dynamics of a system, the P-bifurcation has been deemed too limiting an approach to fully explore the behaviour of a stochastic dynamical system [112, 115]. This is because D-bifurcations have the ability to capture *all* of the stochastic dynamics of a system, while P-bifurcations are limited to the long-run average behaviour, and thus in general may not reflect all of the changes in the stability of the dynamical system. Furthermore, the study of P-bifurcations is largely restricted to dynamical systems driven by white noise. This may be limiting in the study of GRNs, where extrinsic noise is reported to be coloured, with autocorrelation times similar to that of the cell cycle [77]. The dynamical approach may therefore offer a richer understanding of the dynamical structure of a system than the phenomenological approach alone. Indeed, a number of biological and gene regulatory studies have emerged that investigate the dynamical behaviour of a stochastic system from both perspectives [116, 117]. For a comprehensive analysis on P- and D-bifurcations in one-dimensional dynamical systems, we refer the reader to [112], and for an explicit consideration of noise-induced transitions (restricted to P-bifurcations only) we recommend the extensive work of [12].

#### Box 3 Stochastic Bifurcation Theory

##### Bifurcations: deterministic to stochastic

A bifurcation is defined as a qualitative change in the behaviour of a dynamical system in response to varying a parameter, or set of parameters; the bifurcation parameter(s) [109]. Bifurcations are related to the arrival of new solutions and their relative stability. While bifurcation theory has been extensively developed for deterministic systems, relatively little progress has been made towards extending the theory to stochastic dynamical systems.

##### Approaching stochastic bifurcations

Much of the work so far in developing a bifurcation theory for stochastic dynamical systems has been made by Arnold [112], where stochastic bifurcations are distinguished as either “static” (based on the FPE) or “dynamic” (based on the Lyapunov exponents). More specifically, stochastic bifurcations may be viewed in terms of: (i) *Phenomonological*(P)-bifurcations that study qualitative changes in the steady-state probability distribution of a stochastic dynamical system (e.g. from unimodal to bimodal); and (ii) *Dynamical*(D)-bifurcations that study the stability of the dynamical structure of a stochastic system with respect to so-called invariant measures – stochastic analogues of fixed points. The stochastic D-bifurcation is identified as a sign change in the maximal Lyapunov exponent. The two notions are independent of one another and do not necessarily occur together [114, 115].

##### P-bifurcations in developmental models

P-bifurcations have been extensively studied in a number of biological settings with particular application to developmental models. Indeed, key features of the developmental process such as binary cell-fate choices, or lineage commitments, are naturally and frequently viewed in this context. Here the appearance or disappearance of a stable state in the system dynamics can be used to explain the appearance or disappearance of a particular phenotype. Bifurcations thus lend specific insight into the process of cell-fate decision making and commitment, as well the identification of drivers of the underlying system. Traditionally, P-bifurcations have been studied in the context of deterministic dynamical systems. In the presence of noise however, dynamical systems may exhibit rich behaviours. Among these, are bifurcations driven solely by varying the intensity of noise, even as the underlying deterministic system remains unchanged. Such transitions thus lead to the creation or destruction of phenotypes, without changing the parameters of the system. This phenomenon has been observed in a number of developmental contexts, including in the modulation of cellular plasticity [67], cell cycle regulation [89], cancer cell progression and metastases [118], and metabolism [119]. We refer the reader to [2] for a comprehensive overview on the different types of P-bifurcations, noise-induced transitions and their relation to cell-fate decisions.

### Discussion

Here we provide an analytically rigorous, yet intuitive and instructive demonstration of the subtleties that may arise, and the consequences thereof, when computing the landscape in the presence of noise. While some of these complexities may be well-appreciated in certain fields, considering the diverse audience the probabilistic landscape may appeal to, it is important to offer this mathematical framework in a more digestible manner.

We consider how different noise types affect the landscape. In particular, we demonstrate the oft-observed finding that additive noise results only in a disorganising effect on the landscape. In the presence of intrinsic or extrinsic multiplicative noise, the effects may be more profound, resulting in significant deviations between the deterministic and probabilistic landscapes. While these deviations are to some extent noted in the landscape literature [82, 32], we believe that the reason for them is less clear, often buried in a mound of mathematical jargon, rendering the rationale inaccessible to many of us who need it. In short, there is a subtle interplay between the stochastic and deterministic components, with the resultant probabilistic landscape arising as the balance of these influences. It is perhaps unsurprising that significant distortion of the topological features of the landscape can arise when these influences are in conflict. By varying the stochastic component on a fixed deterministic system, we show that the fixed points of the system may be preserved, shifted or destroyed. Sometimes even new fixed points can emerge. It is exactly these changes that are associated with stochastic P-bifurcations. We point out that P-bifurcations may be limited in capturing the full dynamics of the underlying stochastic system and we may therefore additionally want to consider D-bifurcations – the natural analogue of deterministic bifurcations.

A further subtlety arises from the nuanced mathematical treatment of stochasticity in the form of white noise, specifically when it enters the system multiplicatively. We show that the two most commonly employed stochastic integration conventions can produce qualitatively different steady-state distributions and subsequently, different probabilistic landscapes. As these deviations are expected to be appreciable in modelling real-world biological systems, we hope that this emphasises the importance in considering a stochastic differential equation in conjunction with a stochastic calculus when constructing the probabilistic landscape.

While additive noise is often employed in practise [26, 32, 120] – perhaps because it avoids the intricacies of stochastic calculus and the associated consequences discussed herein – it does not satisfy two of the most commonly observed features of noise in cellular differentiation: (i) that the noise is high near the bifurcation point or unstable equilibria, and (ii) is comparatively lower at the differentiated cell states or stable equilibri [67]. A natural, and more realistic noise form that is consistent with these properties is when the noise is proportional to the deterministic landscape itself, or more generally, if it observes a power relationship to the deterministic landscape. We show that in these instances, the fixed points of the underlying deterministic system will be preserved by the probabilistic landscape. Thus, in situations where it is desirable to preserve the dynamical information of the deterministic dynamics, it may not be necessary to turn to alternate methods such as those presented in [82] to compute the landscape, which may be complicated to implement in practice.

An important consideration is the identifiability of the deterministic dynamics in the presence of noise. We show that one-dimensional stochastic systems can be characterised by different deterministic and different stochastic components that give rise to the same stationary distribution. Thus, even if the probabilistic landscape is obtained, we cannot make definitive assumptions about the underlying deterministic and stochastic components of the system on the basis of this information alone. Despite this, non-identifiability may be resolved (at least for one-dimensional systems) by obtaining additional information on any one of the other components of the system, provided we fix *a priori* a stochastic integration convention.

While the examples discussed herein are for one-dimensional gradient systems, we might not be completely discouraged by this since gradient systems are representative of more general dynamical systems. Moreover, for high-dimensional, non-equilibrium systems, the deterministic dynamics can be decomposed into two components, the gradient of a potential and the curl flux. Thus, even in higher dimensions, we are still ultimately concerned with gradient dynamics, which will inevitably be subject to the same mathematical nuances we present here, even before the consequences of curl flux are considered. In [22], for example, the authors consider a two-dimensional, non-equilibrium system and show that increasing levels of additive noise exert only a disorganising effect on the landscape, as we have shown here. Furthermore, there may be cases where a significant reduction in the dimensionality of the data preserves the basic dynamics of the system, allowing us to still gain meaningful insight into the high-dimensional dynamics [36, 121, 122].

We hope that this careful and explicit consideration of noise on the potential landscape will draw the researcher’s attention to the associated complexities and limitations. These are not merely fringe technicalities that can be ignored, but rather significant mathematical caveats with fundamental implications for constructing the landscape as the evolution of a dynamical system in the presence of stochastic fluctuations.

### Methods

#### Derivation of the general steady-state probability distribution

We begin by providing full details of the derivation to Eq. (7) of the main text. The solution is well known, but we include the derivation for the sake of completeness. Consider first the Langevin Equation of a one-dimensional stochastic differential equation (SDE), where the dependence on the stochastic integration convention *α* is explicit,

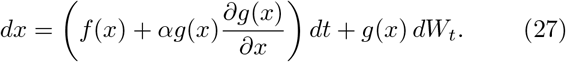

The associated Fokker-Planck equation (FPE) [123] is obtained by inspection of Eq. (27), and can be written as

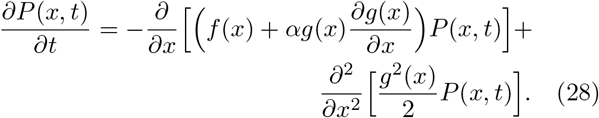

The steady-state solution, denoted *P*_*s*_(*x*), is obtained from Eq. (28) by setting 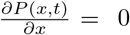. The equation for *P*_*s*_(*x*) may then be written as

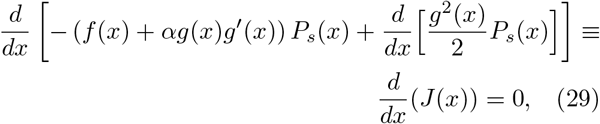

where 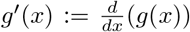. Here *J* denotes the probability flux, and is easily seen from Eq. (29) to be constant. If *J* is 0 everywhere (i.e. the system satisfies the detailed balance condition [123]), then Eq. (29) reduces to

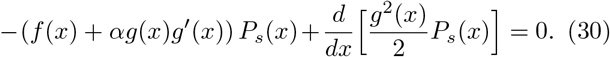

We now make the substitution

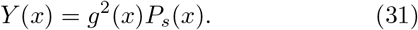

Substituting Eq. (31) into Eq. (30) we obtain the following equation in *Y*,

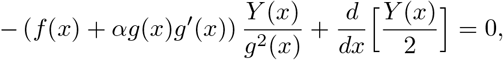

which is simply a first-order linear homogeneous differential equation, and is easily solved as

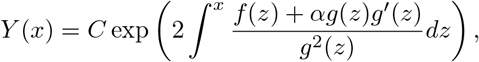

where *C* ∈ ℝ. It follows that the general form of the steady-state solution of the FPE is given by:

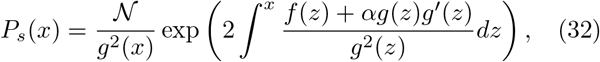

where 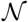 is the normalisation constant. This is Eq. (7) in the main text. It then follows immediately, from Eq. (4) in the main text, that the probabilistic landscape *U*_*q*_(*x*) (Eq. (8) in the main text), is given by

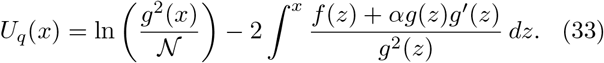

#### Example 1: Shifting fixed points

In the main text, the first example of multiplicative noise has the noise component 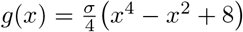. Here we provide details of the derivation of the stationary probability distribution, and corresponding probabilistic landscape. From the general solution to the steady-state probability distribution *P*_*s*_(*x*) (Eq. (7) in the main text), with *f*(*x*) = *x* − *x*^3^ and 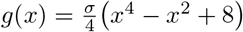, we have

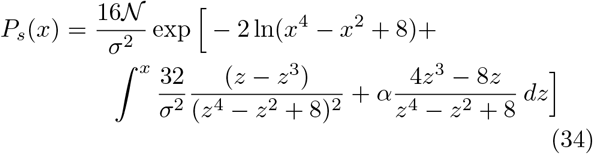

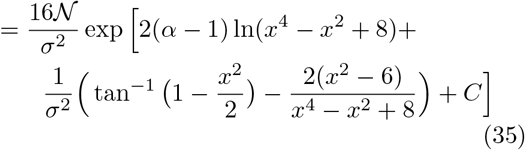

where 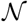 is the normalisation constant and *C* is the constant of integration. It then follows immediately from Eq. (8) in the main text that

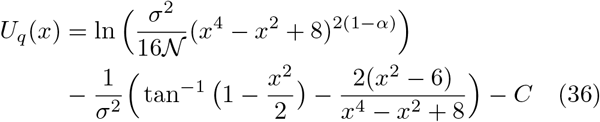

In Fig. 3 in the main text, the value for *C* is taken to be 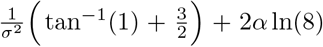, and the normalisation constant is calculated accordingly.

In the main text, we discussed an additional example that exhibits behaviour similar to that of Example 1, except now the fixed points of the system are shifted towards the origin (as opposed to being pushed away from the origin as in Example 1). We provide here details of this example. If we consider the noise component 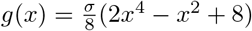 on the double-well potential *U*(*x*) defined in Eq. (9) of the main text, then it can be shown that the steady-state probability distribution of the state of the system *x* is given by

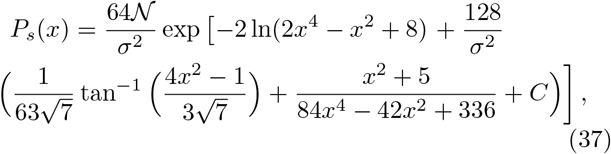

where 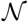 is the normalisation constant and *C* is the constant of integration. In Fig. S3(A), we plot the effects of the noise *g*(*x*) (for *σ* = 0.5) on the double-well potential *U*(*x*). We can see from Fig. S3(B&C) that as the noise intensity *σ* increases, the fixed points of steady-state distribution *P*_*s*_(*x*), and corresponding probabilistic landscape *U*_*q*_(*x*), shift towards those of the noise curve *g*(*x*), which occur at 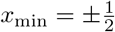.

#### Example 2: Destruction of fixed points

We provide full details of the derivations to the stationary probability distribution and probabilistic landscape given in Example 2 of the main text. We also derive the stationary points of the probabilistic landscape, which we use to calculate the critical noise, *σ*_*c,α*_, at which the stochastic system transitions from bistable to monostable.

#### Derivation of the steady-state distribution and probabilistic landscape

From the general solution of the steady-state probability distribution (Eq. (7)), with *f*(*x*) = *x − x*^3^ and *g*(*x*) = *σ*(1 + *x*^2^) (*σ >* 0), we have

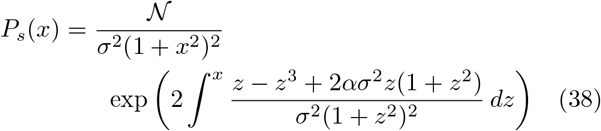

where 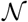 is the normalisation constant. Consider first the integral in Eq. (38) which, after using partial fractions, can be written as

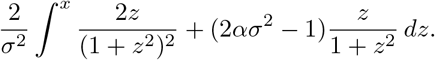

Now, letting *u* = *z*^2^ so that 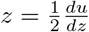, we obtain

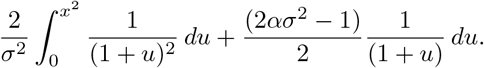

Then integrating with respect to *u* we obtain,

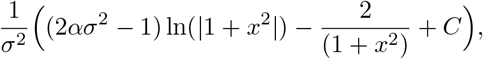

where *C* is the constant of integration. It follows that

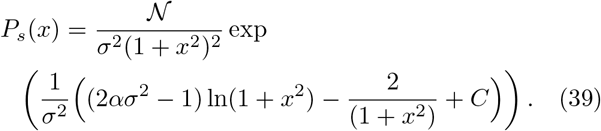

This is Eq. (14) in the main text. Now computing the probabilistic landscape, we immediately obtain

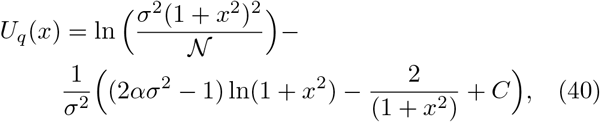

which is Eq. (15) in the main text. For Fig. 4 in the main text we choose *C* = 2, and then calculate the normalisation constant accordingly.

#### Stable stationary solutions of the probabilistic landscape

Differentiating *U*_*q*_(*x*) (as given by Eq. (40) above) with respect to *x* and setting *U′*_*q*_(*x*)=0, we obtain (after simplification),

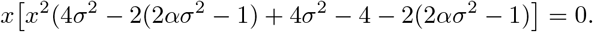

This implies that either *x* = 0 or

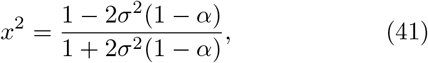

which gives rise to Eq. (16) in the main text.

#### The Schlögl reaction network

We consider here the effect of an additive noise term on the dynamics of the Schlögl reaction network. The purpose of this exploration is to compare with the analysis presented in the main text, where we consider the effects of multiplicative “Langevin” noise. For these results, along with a discussion of the comparison, refer to the section titled “Effect of coupled noise on the landscape” in the main text. In Fig. S1(A), we display both the potential function *U*(*x*) of the Schlögl network (Eq. (20) of the main text) and additive noise function *g*(*x*) = *σ*(plotted for *σ* = 0.1); these are the purple and red curves, respectively. The deterministic potential function is plotted for parameters *k*_1_ = 0.019, *k*_2_ = 0.25, *k*_3_ = 1, and *k*_4_ = 1.18. For ease of visualisation, we have chosen different parameters to those considered in the main text, but emphasise that this does not change the conclusions of our comparison: the effect of additive noise on the deterministic landscape is independent of the specific potential function *U*(*x*). As expected from Eq. (11) of the main text, additive noise does not lead to qualitative changes in the landscape: the effect is only to scale and (vertically) shift the potential; as demonstrated in Fig. S1(B&C).

#### Preserving fixed points under multiplicative noise

In the main text, we claim that if the noise component *g*(*x*) is related to the potential landscape *U*(*x*) by way of the following power relationship,

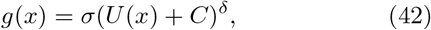

where *σ >* 0 is the noise intensity and *C, δ* are any non-negative real numbers, then provided that *U*(*x*) + *C* is strictly positive, the fixed points of *U*_*q*_(*x*) will correspond precisely to those of *U*(*x*), when they exist. Here we provide a proof of this result. We first remind the reader that since we are working with gradient systems, the deterministic component *f*(*x*) is equal to the negative gradient of the potential *U*(*x*) (see Eq. (1) of the main text). The proof proceeds in two cases: (1) *α* ∈ [0, 1) and (2) *α* = 1. We begin by assuming that *α* ∈ [0, 1), then from the general solution for *P*_*s*_(*x*) (Eq. (7)) we have,

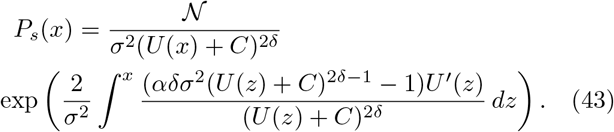

Now letting *v* = *U*(*x*) + *C* so that *v*′ = *U*′(*x*), the integral in Eq. (43) becomes

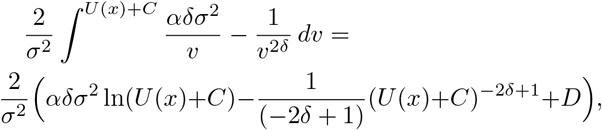

for 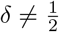, and

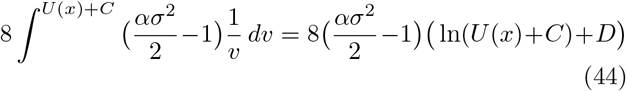

for 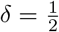, where *D* is some constant.

When 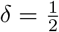, it is easy to show that *P*′_*s*_(*x*) = 0 is equivalent to *U*′(*x*) = 0 (this follows immediately after calculating *P*′_*s*_(*x*)). We now consider the case where 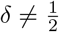. In this case, Eq. (43) becomes,

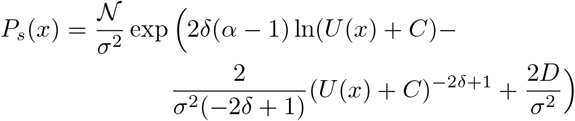

Then calculating *P*_*s*_′(*x*), we have that *P*_*s*_′(*x*) = 0 is equivalent to

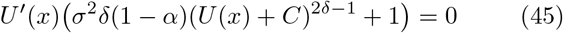

If *δ* = 0, this reduces to *U*′(*x*) = 0, and we are done. Now, assume that *δ >* 0. It then follows from Eq. (45) that either *U*′(*x*) = 0 or

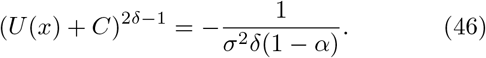

As *α* ∈ [0, 1) we have that (1 − *α*) ∈ (0, 1], and since *δ >* 0, the right hand side of Eq. (46) is negative. Now, as *U*(*x*) + *C* is strictly positive by assumption, it follows that Eq. (46) has no solutions. Thus, the stationary points of *P*_*s*_(*x*), and therefore of *U*_*q*_(*x*), will agree with those of *U*(*x*) when they exist. The case for *α* = 1 is similar. The proof proceeds in the same way as for Case (1), except that now Eq. (45) reduces to *U*′(*x*) = 0, and so we immediately obtain the desired result.

We now briefly consider an example of noise satisfying the power relationship in Eq. (42) acting on the double-well potential *U*(*x*) defined in Eq. eq:model of the main text. It can be shown that the steady-state probability distribution for the state of the system *x* is given by

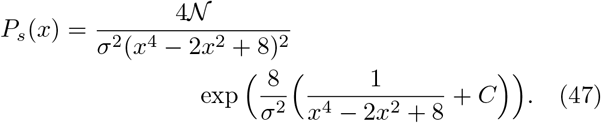

In Fig. S2(A), we display the effects of the noise *g*(*x*) = *σ*(*U*(*x*) + 2) on the double-well potential *U*(*x*). In Fig. S2(B&C), we display the steady-state distribution and corresponding landscape for the Itô interpretation (the Stratonovich interpretation produces very similar results). We can see that the effect of noise is qualitatively indistinguishable from that of additive noise (compare with Fig. 2 in the main text).

#### Decomposing the stationary probability distribution

The stationary probability distribution *P*_*s*_(*x*) of the SDE given in Eq. (27) is governed by the FPE given in Eq. (29), and can be seen to reduce to the differential equation given in Eq. (30), provided the detailed balance condition holds. We can use the differential equation given in Eq. (30) to *uniquely* solve for any one of the components *f*(*x*), *g*(*x*) or *P*_*s*_(*x*) in terms of the other two; the uniqueness follows from the uniqueness property of first-order differential equations [103].

We begin by fixing a stochastic integration convention *α*. Then given any stationary probability distribution *P*_*s*_(*x*) and noise component *g*(*x*), the deterministic component *f*(*x*) can be easily solved by rearranging the differential equation Eq. (30). We obtain

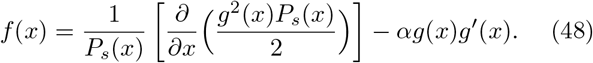

This is Eq. (24) in the main text. Now, given any stationary probability distribution *P*_*s*_(*x*) and deterministic component *f*(*x*), the stochastic component *g*(*x*) can be solved from the differential equation Eq. (30) as,

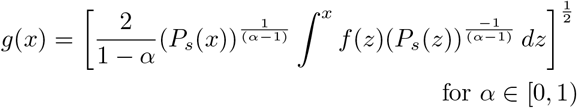

and

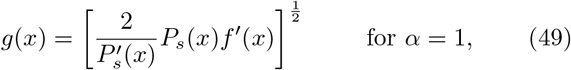

which is Eq. (25) in the main text. We now check that this is the solution to differential equation given in Eq. (30). It easy to see that Eq. (30) is equivalent to

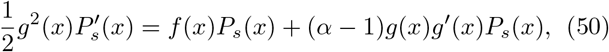

We consider two cases. First consider the case where *α* = 1. It follows immediately from Eq. (50) that

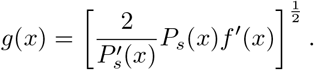

Now assume that *α* ∈ [0, 1). Then from Eq. (50) we have

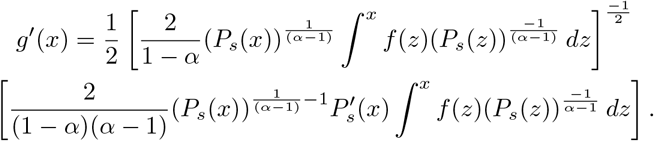

So that,

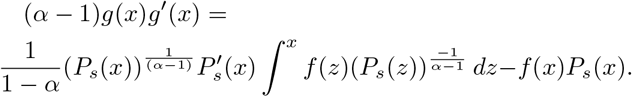

Thus, the right hand side of Eq. (50) is equal to

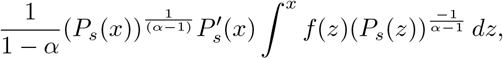

which equals the left hand side of Eq. (50). Hence *g*(*x*) is indeed the solution to differential equation given in Eq. (30).

#### Non-identifiability of the first kind

Here we derive the stationary probability distribution *P*_*s*_(*x*) (Eq. (26) of the main text) and deterministic component 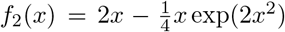 used in the main text to demonstrate non-identifiabilty of the first kind. We begin by deriving *P*_*s*_(*x*). From the general solution of the steady-state probability distribution (Eq. (7) of the main text), with *f*_1_(*x*) = −*x* and *g*_1_(*x*) = *σ* exp(−*x*^2^), we have

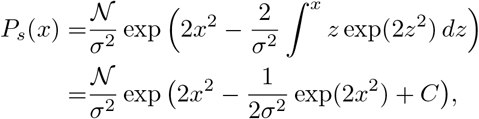

where 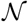 is the normalisation constant and *C* is the constant of integration. Now, using Eq. (48), with *g*_2_(*x*) = 1 and *P*_*s*_(*x*), the deterministic component can be solved as

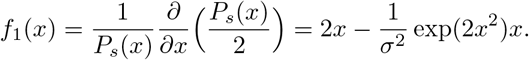

## Author contributions

M.A.C., L.H. and M.P.H.S. conceptualised the research. Formal analysis was conducted by M.A.C. and L.H. Writing by M.A.C. and L.H., with editing and review contributed by M.P.H.S. All authors provided critical feedback and helped shape the research.

## Acknowledgments

We gratefully acknowledge Rowan D. Brackston for helpful discussions in the early stages of this research, as well the support from the members of the *Theoretical Systems Biology Group* at the University of Melbourne. L.H. and M.P.H.S. are supported by the University of Melbourne DVCR fund. M.A.C is supported by the University of Melbourne graduate research scholarship.

## Supplementary material

### Derivation of the general steady-state probability distribution

We begin by providing full details of the derivation to Eq. (6) of the main text. The solution is well known, but we include the derivation for the sake of completeness. Consider first the Langevin Equation of a one-dimensional stochastic differential equation (SDE), where the dependence on the stochastic integration convention *α* is explicit,

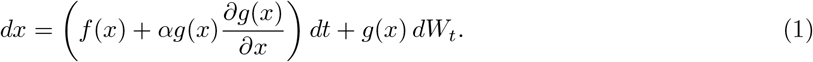

The associated Fokker-Planck equation (FPE) [1] is obtained by inspection of (1), and can be written as

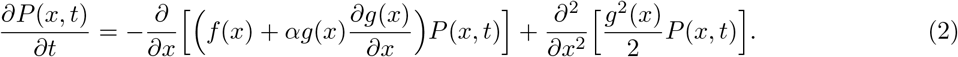

The steady-state solution, denoted *P*_*s*_(*x*), is obtained from Eq. (2) by setting 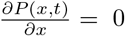. The equation for *P*_*s*_(*x*) may then be written as

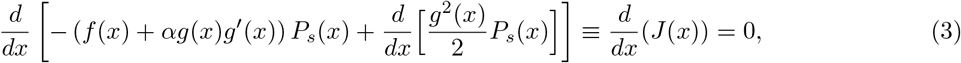

where 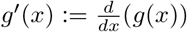. Here *J* denotes the probability flux, and is easily seen from (3) to be constant. If *J* is 0 everywhere (i.e. the system satisfies the detailed balance condition [1]), then Eq. (3) reduces to

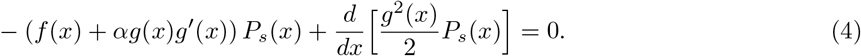

We now make the substitution

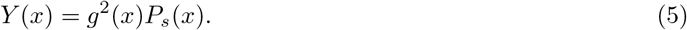

Substituting Eq. (5) into Eq. (4) we obtain the following equation in *Y*,

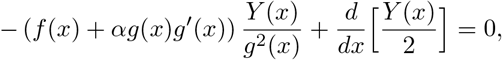

which is simply a first-order linear homogeneous differential equation, and is easily solved as

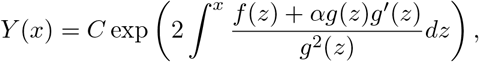

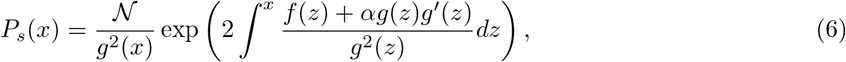

where 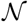 is the normalisation constant. This is Eq. (6) in the main text. It then follows immediately, from Eq. (3) in the main text, that the probabilistic landscape *U*_*q*_(*x*) (Eq. (7) in the main text), is given by

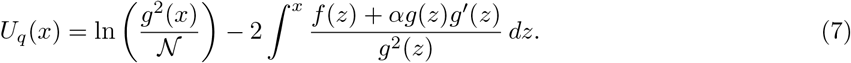

**Figure 1:**
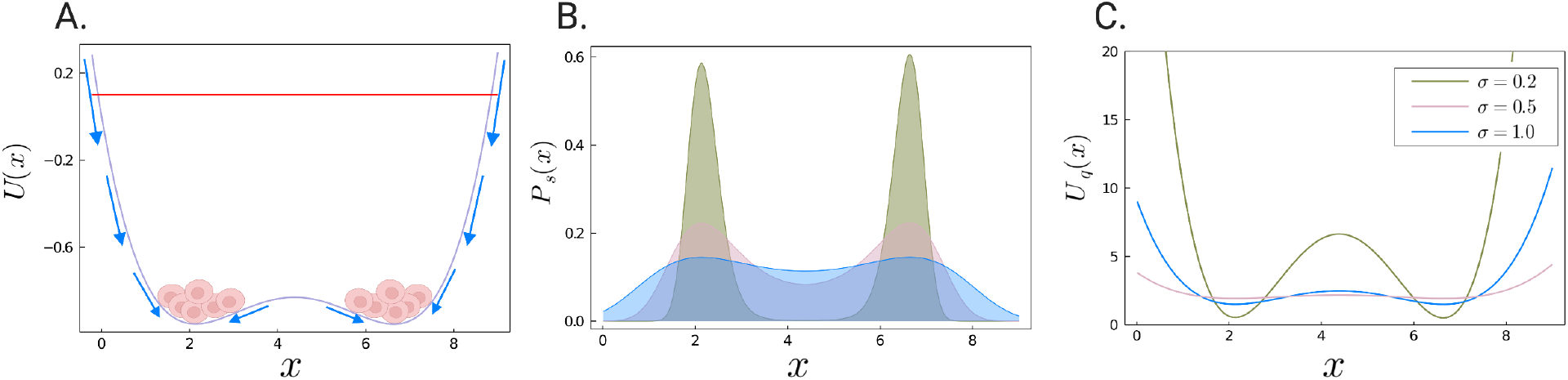
The effect of additive noise on the Schlögl reaction network. (A) A schematic demonstrating the effect of additive noise on the potential landscape; details of the parameters are given in the text. (B) Analytical steady-state probability distribution *P*_*s*_(*x*) given by Eq. (22) for *k*_1_ = 0.019, *k*_2_ = 0.25, *k*_3_ = 1, *k*_4_ = 1.18. (C) The corresponding probabilistic landscape *U*_*q*_(*x*). Both figures are plotted for increasing strengths of additive noise. The different colours represent the different levels of noise: the green curve is for *σ* = 0.2, the pink curve for *σ* = 0.5 and the blue curve for *σ* = 1.0. Increasing the noise parameter *σ* flattens both *P*_*s*_(*x*) and *U*_*q*_(*x*), making rare transitions between states more probable, however the positions of the maxima of *P*_*s*_(*x*) and minima of *U*_*q*_(*x*) are preserved.

### Multiplicative noise

#### The Schlögl reaction network

We consider here the effect of an additive noise term on the dynamics of the Schlögl reaction network. The purpose of this exploration is to compare it with the analysis presented in the main text, where we consider the (multiplicative) “Langevin’ noise. For these results, along with a discussion of the comparison, refer to the section “Effect of coupled noise on the landscape” in the main text. In Fig. 1(A), we display both the potential function *U*(*x*) of the Schlögl network (Eq. (20) of the main text) and additive noise function *g*(*x*) = *σ*(plotted for *σ* = 0.1); these are the purple and red curves, respectively. The deterministic potential function is plotted for parameters *k*_1_ = 0.019, *k*_2_ = 0.25, *k*_3_ = 1, and *k*_4_ = 1.18. For ease of visualisation, we have chosen different parameters to those considered in the main text, but emphasise that this does not change the conclusions of our comparison: the effect of additive noise on the deterministic landscape is independent of the specific potential function *U*(*x*). As expected from Eq. (11) of the main text, additive noise does not lead to qualitative changes in the landscape: the effect is only to scale and (vertically) shift the potential; as demonstrated in Fig. 1(B&C) above.

#### Preserving fixed points under multiplicative noise

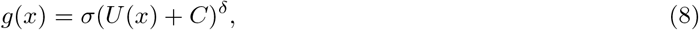

where *σ >* 0 is the noise intensity and *C, δ* are any non-negative real numbers, then provided that *U*(*x*) + *C* is strictly positive, the fixed points of *U*_*q*_(*x*) will correspond precisely to those of *U*(*x*), when they exist. Here we provide a proof of this result. First we remind the reader that since we are working with gradient systems, the deterministic component *f*(*x*) is equal to the negative gradient of the potential *U*(*x*) (see Eq. (1) of the main text). The proof proceeds in two cases: (1) *α* ∈ [0, 1) and (2) *α* = 1. We begin by assuming that *α* ∈ [0, 1), then from the general solution for *P*_*s*_(*x*) (Eq. (6)) we have,

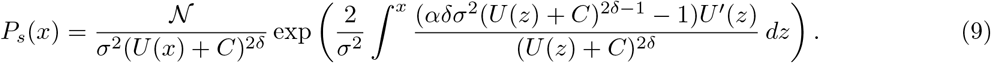

**Figure 2:**
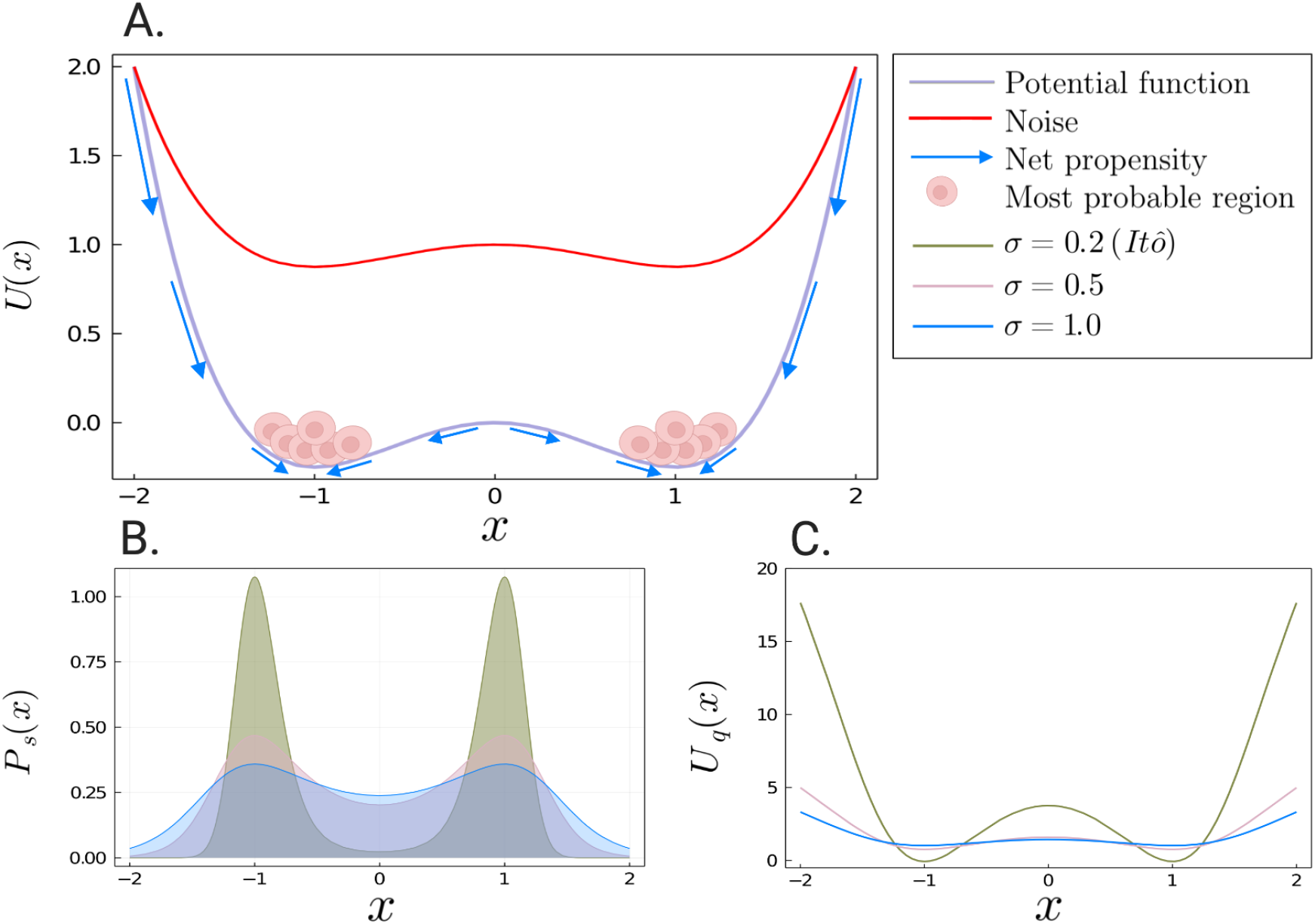
(A) A schematic of the system’s trajectory. Here we consider a deterministic double-well potential *U*(*x*) driven by multiplicative noise satisfying the power relationship given in Eq. (8) with *C* = 2 and *δ* = 1. (B&C) Analytical steady-state probability distribution *P*_*s*_(*x*) for the Ito interpretation *α* = 0. We consider the effects of increasing strengths of multiplicative noise *g*(*x*) = *σ*(*U*(*x*) + 2), where *σ* = 0.2 (green curve), 0.5 (pink curve), 1.0 (blue curve). As *σ* increases, the maxima of *P*_*s*_(*x*) and minima of *U*_*q*_(*x*) remain fixed.

Now letting *v* = *U*(*x*) + *C* so that *v*′ = *U*′(*x*), the integral in Eq. (9) becomes

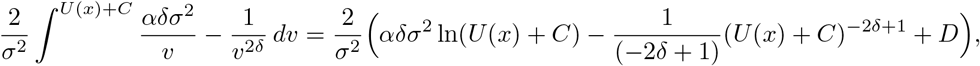

for 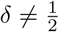, and

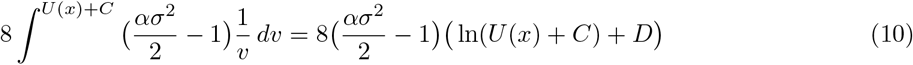

for 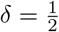, where *D* is some constant.

When 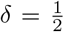, calculating *P*′_*s*_(*x*) it is easy to show that *P*′_*s*_(*x*) = 0 is equivalent to *U*′(*x*) = 0. We now consider the case where 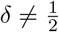. Then Eq. (9) becomes,

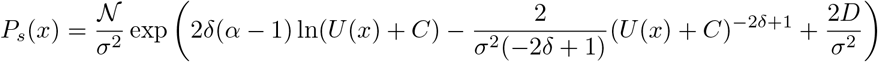

Then calculating *P*_*s*_′(*x*), we have that *P*_*s*_′(*x*) = 0 is equivalent to

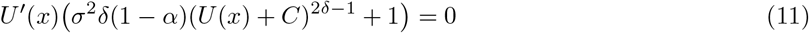

If *δ* = 0, this reduces to *U*′(*x*) = 0, and we are done. Now, assume that *δ >* 0. It then follows from Eq. (11) that either *U*′(*x*) = 0 or

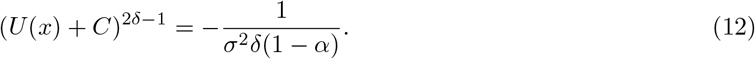

As *α* ∈ [0, 1) we have that (1 − *α*) ∈ (0, 1], and since *δ >* 0, the right hand side of Eq. (12) is negative. Now, as *U*(*x*) + *C* is strictly positive by assumption, it follows that Eq. (12) has no solutions. Thus, the stationary points of *P*_*s*_(*x*), and therefore of *U*_*q*_(*x*), will agree with those of *U*(*x*) when they exist. The case for *α* = 1 is similar. The proof proceeds in the same way as for Case (1), except that now Eq. (11) reduces to *U*′(*x*) = 0, and so we immediately obtain the desired result.

We now briefly consider an example of noise satisfying the power relationship in Eq. (8) acting on the double-well potential *U*(*x*) (defined in Eq. (8) of the main text). It can be shown that the steady-state probability distribution for the state of the system *x* is given by

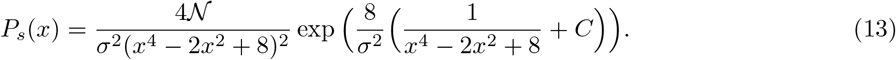

In Fig. 2(A), we display the effects of the noise *g*(*x*) = *σ*(*U*(*x*) + 2) on the double-well potential *U*(*x*). In Fig. 2(B&C), we display the steady-state distribution and corresponding landscape for the Itô interpretation (the Stratonovich interpretation produces very similar results). We can see that the effect of noise is qualitatively indistinguishable from that of additive noise (compare with Fig. 2 of the main text).

#### Example 1: Shifting fixed points

In the main text, the first example of multiplicative noise is for the noise component 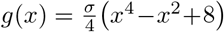. Here we provide details of the derivation of the stationary probability distribution, and corresponding probabilistic landscape. From the general solution to the steady-state probability distribution *P*_*s*_(*x*) (Eq. (6) in the main text), with *f*(*x*) = *x* − *x*^3^ and 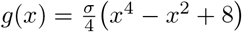, we have

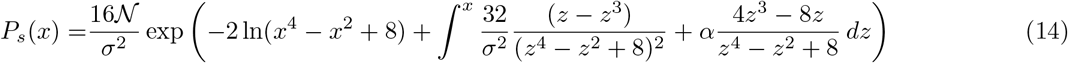

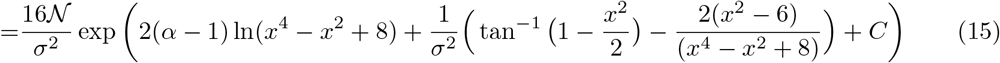

where 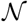 is the normalisation constant and *C* is the constant of integration. It then follows immediately from Eq. (7) in the main text, that

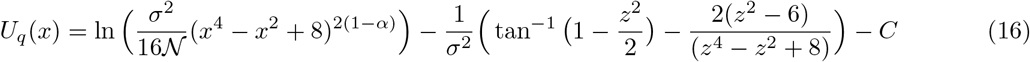

In Fig. (3) in the main text, the value for *C* is 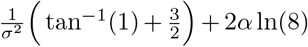, and the normalisation constant is calculated accordingly.

In the main text, we discussed an additional example that behaves in a similar way to that of Example 1, but shifts the fixed points of the system towards the origin (as opposed to being pushed away from the origin as in Example 1). We provide here details of this example. If we consider the noise component 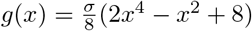 on the double-well potential *U*(*x*) (defined in Eq. (8) of the main text), then it can be shown that the steady-state probability distribution of the state of the system *x* is given by

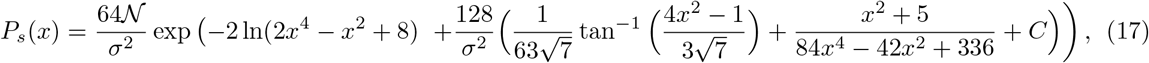

where 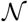 is the normalisation constant and *C* is the constant of integration. In Fig. 3(A), we plot the effects of the noise *g*(*x*) (for *σ* = 0.5) on the double-well potential *U*(*x*). We can see from Fig. 3(B&C) that as the noise intensity *σ* increases, the fixed points of steady-state distribution *P*_*s*_(*x*) and corresponding probabilistic landscape *U*_*q*_(*x*), shift towards those of the noise curve *g*(*x*), which occur at 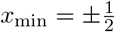).

**Figure 3:**
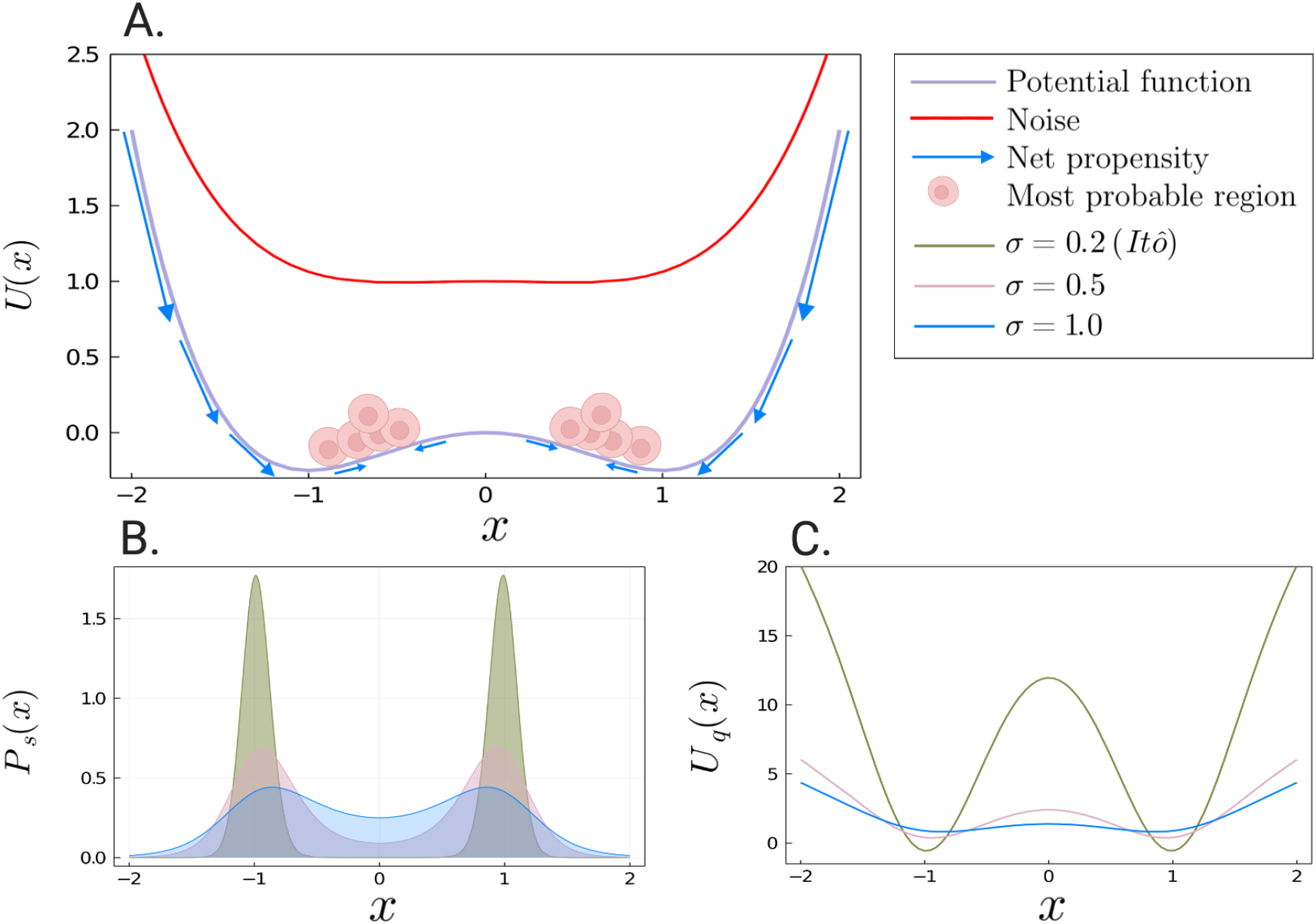
(A) A schematic of the system’s trajectory. We consider the deterministic double-well potential defined in Eq. (8) of the main text driven by multiplicative noise. The schematic is plotted for *σ* = 0.5 Cells are driven *towards* the fixed points of the noise curve 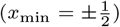. (B&C) Analytical steady-state probability distribution *P*_*s*_(*x*) given by Eq. (17) for the It0̂ interpretation only. We consider the effects of increasing strengths of multiplicative noise 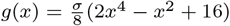, where *σ* = 0.2 (green curve), 0.5 (pink curve), 1.0 (blue curve). As *σ* increases, the maxima of *P*_*s*_(*x*) and minima of *U*_*q*_(*x*) drift towards origin, approaching 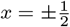.

#### Example 2: Destruction of fixed points

We provide full details of the derivations to the stationary probability distribution and probabilistic landscape given in Example 2 of the main text. We also derive the stationary points of the probabilistic landscape, which we use to calculate the critical noise at which the stochastic system transitions from bistable to monostable.

#### Derivation of the steady-state distribution and probabilistic landscape

From the general solution of the steady-state probability distribution (Eq. (6)), with *f*(*x*) = *x − x*^3^ and *g*(*x*) = *σ*(1 + *x*^2^) (*σ >* 0), we have

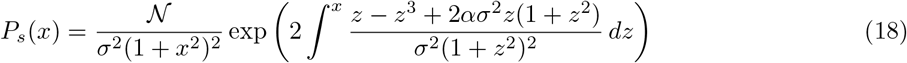

where 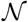 is the normalisation constant. We consider the integral in Eq. (18) which, after using partial fractions, can be written as

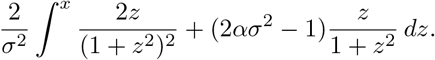

Now, letting *u* = *z*^2^ so that 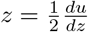, we obtain

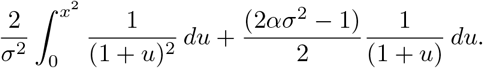

Then integrating with respect to *u* we obtain,

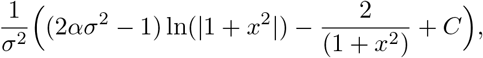

where *C* is the constant of integration. It follows that

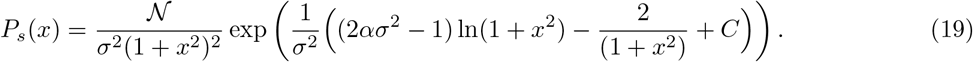

This is Eq. (13) in the main text. Now computing the probabilistic landscape, we immediately obtain

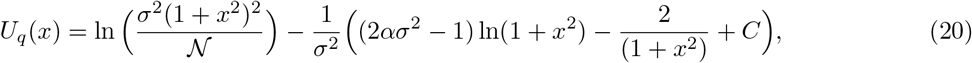

which is Eq. (14) in the main text. Note that for Fig. 4 in the main text we choose *C* = 2, and the normalisation constant is calculated accordingly.

#### Stable stationary solutions of the probabilistic landscape

Differentiating *U*_*q*_(*x*) (as given by Eq. (20) above) with respect to *x* and setting *U′*_q_(*x*)=0, we obtain (after simplification),

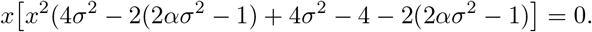

This implies that either x = 0 or

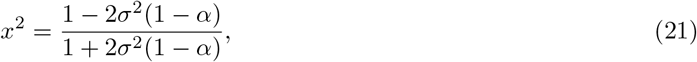

which gives rise to Eq. (19) in the main text.

### Identifiability considerations

#### Decomposing the stationary probability distribution

The stationary probability distribution *P*_*s*_(*x*) of the SDE given in Eq. (1) is governed by the FPE given in Eq. (3), and can be seen to reduce to the differential equation given in Eq. (4), provided the detailed balance condition holds. We can use the differential equation given in Eq. Eq. (4) to *uniquely* solve for any one of the components *f*(*x*), *g*(*x*) or *P*_*s*_(*x*) in terms of the other two; the uniqueness follows from the uniqueness property of first-order differential equations [2].

We begin by fixing a stochastic integration convention *α*. Then given any stationary probability distribution *P*_*s*_(*x*) and noise component *g*(*x*), the deterministic component *f*(*x*) can be easily solved by rearranging the differential equation Eq. (4). We obtain

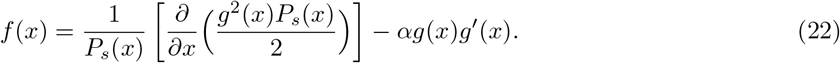

This is Eq. (18) in the main text. Now, given any stationary probability distribution *P*_*s*_(*x*) and deterministic component *f*(*x*), the stochastic component *g*(*x*) can be solved from the differential equation Eq. (4) as,

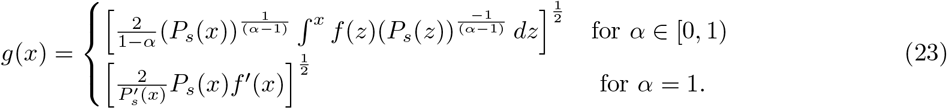

which is Eq. (19) in the main text. We now check that this is the solution to differential equation given in Eq. (4). It easy to see that Eq. (4) is equivalent to

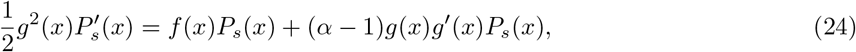

We consider two cases. First consider the case where *α* = 1. It follows immediately from Eq. (23), that

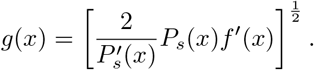

Now assume that *α* ∈ [0, 1). Then from Eq. (23), we have

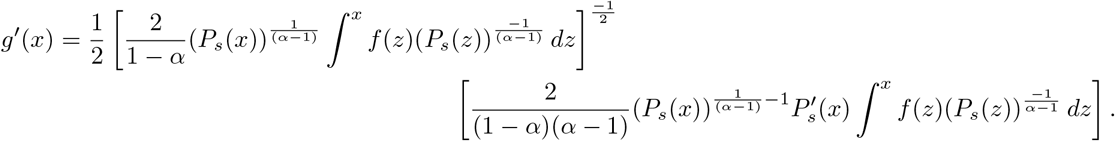

So that,

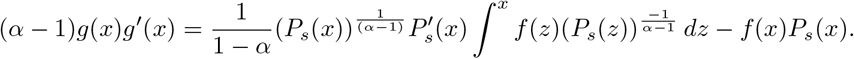

Thus, the right hand side of Eq. (24) is equal to

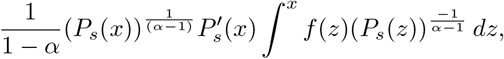

which equals the left hand side of Eq. (24). Hence *g*(*x*) is indeed the solution to differential equation given in Eq. (4).

#### Non-identifiability of the first kind

Here we derive the stationary probability distribution *P*_*s*_(*x*) (Eq. (20) of the main text) and deterministic component 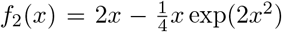 used in the main text to demonstrate non-identifiabilty of the first kind. We begin by deriving *P*_*s*_(*x*). From Eq. (6), with *f*_1_(*x*) = −*x* and *g*_1_(*x*) = *σ* exp(−*x*^2^), we have

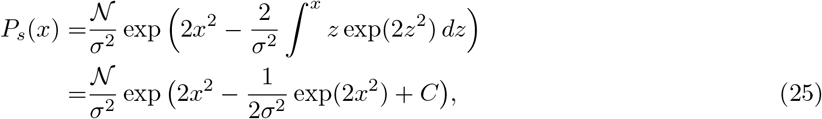

where 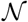 is the normalisation constant and *C* is the constant of integration. Now, using Eq. (22), with *g*_2_(*x*) = 1 and *P*_*s*_(*x*), the deterministic component can be solved as

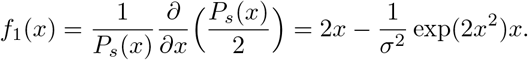

## Footnotes

2 We adopt the terminology of [2], and use *cell states* to describe the transcriptional output of a GRN. Cells progress through a series of cell states to arrive at their eventual cell fate.

3 We adopt the terminology of [2], and use *topology* to describe key features of the landscape such as the number and position of fixed points.

